# scDeepSim: Deep Generative Simulation for Controlled Benchmarking of Single-Cell Methods

**DOI:** 10.64898/2026.11.09.765456

**Authors:** Langtian Ma, Jingyi Jessica Li, Kris Sankaran

## Abstract

Benchmarking single-cell computational methods requires both realistic data and reliable ground truth. However, annotating real datasets is costly and, even when available, may not fully represent the diverse settings where methods are used. Simulators address this gap by generating synthetic data that has fully specified ground truth and whose structure can be tailored to challenge competing methods. Existing single-cell simulators can broadly be divided into *statistical* and *deep generative* classes. Statistical simulators offer interpretable parameters for controlling simulated data, but their modeling assumptions may not accurately represent real data. Deep generative models can learn complex expression patterns, but it remains unclear how to transparently control individual biological or technical factors without introducing unintended effects. We introduce scDeepSim, a simulator that combines a structured variational autoencoder with latent diffusion to generate synthetic single-cell expression profiles. It allows users to intervene on selected biological or technical factors at user-specified strengths, producing realistic datasets with benchmark ground truth. We demonstrate this capability by benchmarking batch correction and trajectory inference methods, showing how batch-effect strength affects batch mixing and cell-type preservation, and how trajectory geometry and noise affect pseudotime recovery. scDeepSim therefore enables researchers to vary selected biological and technical factors in realistic simulated data and assess how these changes affect the performance of single-cell analysis methods, helping understand when such methods may be appropriately applied. Code for the scDeepSim simulator and experiments is available at https://github.com/LangtianM/scDeepSim.

## 1 Introduction

Simulation is an essential device for navigating the landscape of single-cell analysis methods. Simulated data can serve as ground truth for benchmarking methods when annotating real data is costly. Further, while real data are fixed, simulators provide a laboratory for studying how specific data properties affect method performance. For example, simulators can reveal settings where batch effect correction methods overintegrate the data, masking real biological differences [31, 56], and they can rank trajectory inference methods under different assumed topologies [40]. While simulation-based benchmarks are often useful for these reasons, they depend critically on whether the generated data are realistic and whether data properties can be varied in ways that highlight meaningful differences across methods [9]. Therefore, a useful simulator should provide two complementary capabilities: (i) the ability to generate realistic single-cell data and (ii) the ability to vary selected biological and technical factors while preserving realism. Existing simulation approaches provide different ways to support these goals, but there is a basic tension between them.

Mechanistic and statistical simulators provide explicit ways to vary specific factors through user-provided regulatory dynamics or interpretable model parameters [7, 55, 45]. For example, Splatter’s Splat model provides parameters for cell groups, batch effects, and differentiation paths [55]. scDesign3 learns expression distributions and gene dependencies conditional on cell types, batches, and other covariates; modifying its fitted parameters allows selected effects to be varied [45]. These approaches make benchmark conditions interpretable, but the realism of the generated data depends on how well the chosen model captures the data population, and preserving heterogeneity within a labeled cell population remained a challenge [9].

Deep generative models offer a potentially more flexible approach to learning gene expression distributions from observed data [33, 32, 37]. However, they do not offer as straightforward control of the biological or technical factors relevant for benchmarking. For example, scDiffusion and scLDM generate cells from learned latent distributions [32, 37]. Conditional generation allows scDiffusion to sample specific cell populations and interpolate between them [32]. However, the less transparent nature of deep learning-based conditional generation means that varying these factors can have unintended consequences for factors that are not meant to be varied (Section 3.2). A related line of work uses latent representations to separate biological and technical effects for expression prediction [29, 57, 27, 20]. For example, CPA separates perturbation effects from cellular state to predict responses to perturbation combinations [27], and scDisInFact disentangles the effects of different conditions on gene expression [57]. TarDis allocates categorical and continuous covariates to separate latent subspaces to support data integration and counterfactual prediction [20]. These approaches motivate the use of structured representations for controlled benchmarking, where a complementary goal is to sample new cell populations and systematically vary selected factors across matched simulated datasets.

To support this goal, we propose scDeepSim, a deep generative simulator that learns complex single-cell expression patterns while allowing users to reliably control selected biological and technical properties in the generated data (Fig. 1). First, a structured variational autoencoder maps each expression profile to a latent representation with subvectors assigned to different factors. Training encourages each subvector to represent its assigned factor. For example, at full batch intervention strength (*α* = 1) on the scIB human pancreas dataset [31], scDeepSim better preserves cell-type structure, with a mean cell-type average silhouette width (ASW) of 0.519, compared with 0.432 for a plain VAE (Section 3.2). Second, a latent diffusion model learns the distribution of the encoded cells and samples new latent representations. Third, a post-hoc intervention modifies only the subvectors assigned to the target factor while retaining the remaining parts of those representations. Decoding the manipulated representations produces matched datasets whose intervention values can serve as ground truth. We make three contributions:

**Fig. 1.**
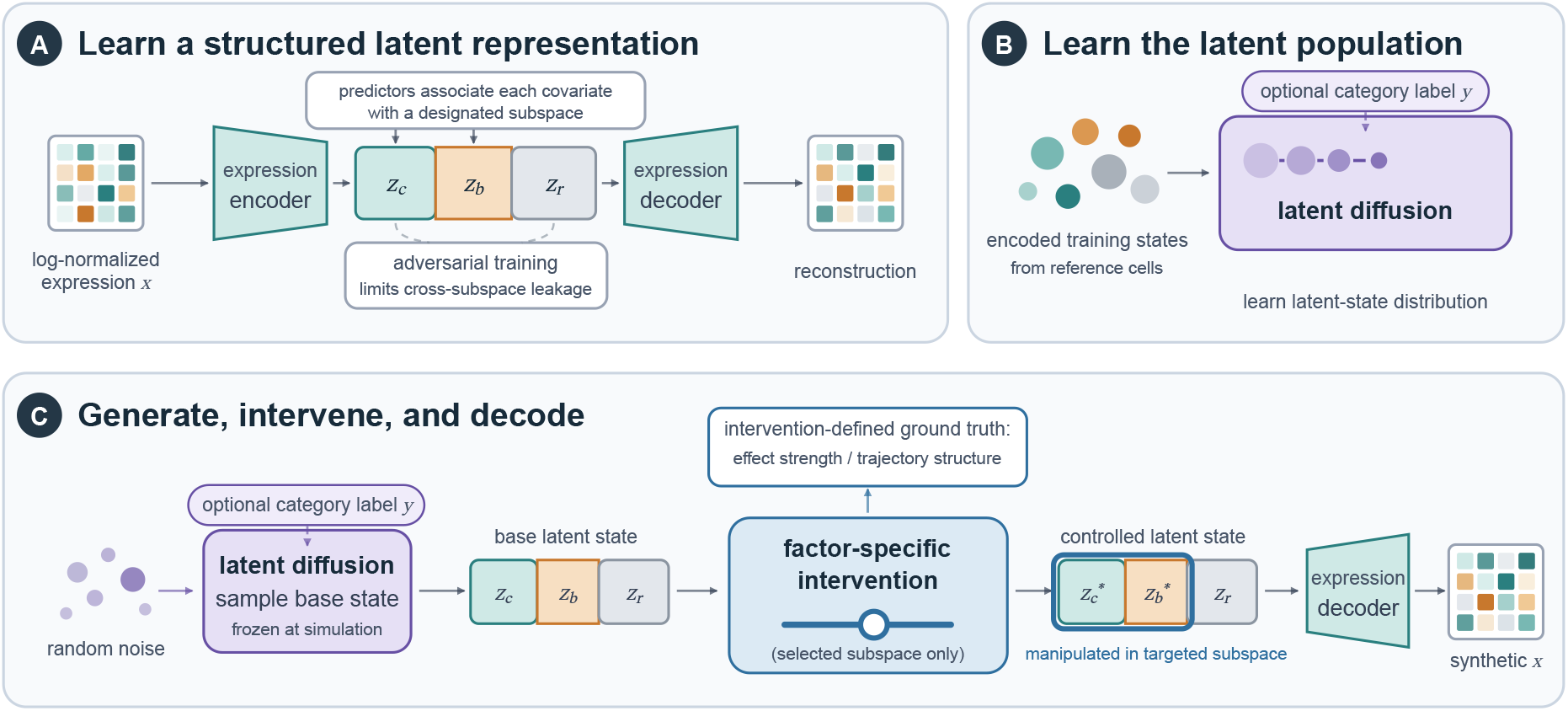
Overview of scDeepSim. (A) A structured variational autoencoder maps an expression profile to latent representations composed of subvectors representing cell type **z**_*c*_, batch **z**_*b*_, and residual variation **z**_*r*_. Supervised and adversarial training encourage each subvector to mainly represent its assigned factor. (B) A latent diffusion model learns the distribution of encoded training cells and samples latent vectors. (C) A post-hoc intervention changes only the selected latent subvector. Decoding the manipulated latent vector yields intervened synthetic datasets, while the intervention parameter provides benchmark ground truth.

**Fig. 2.**
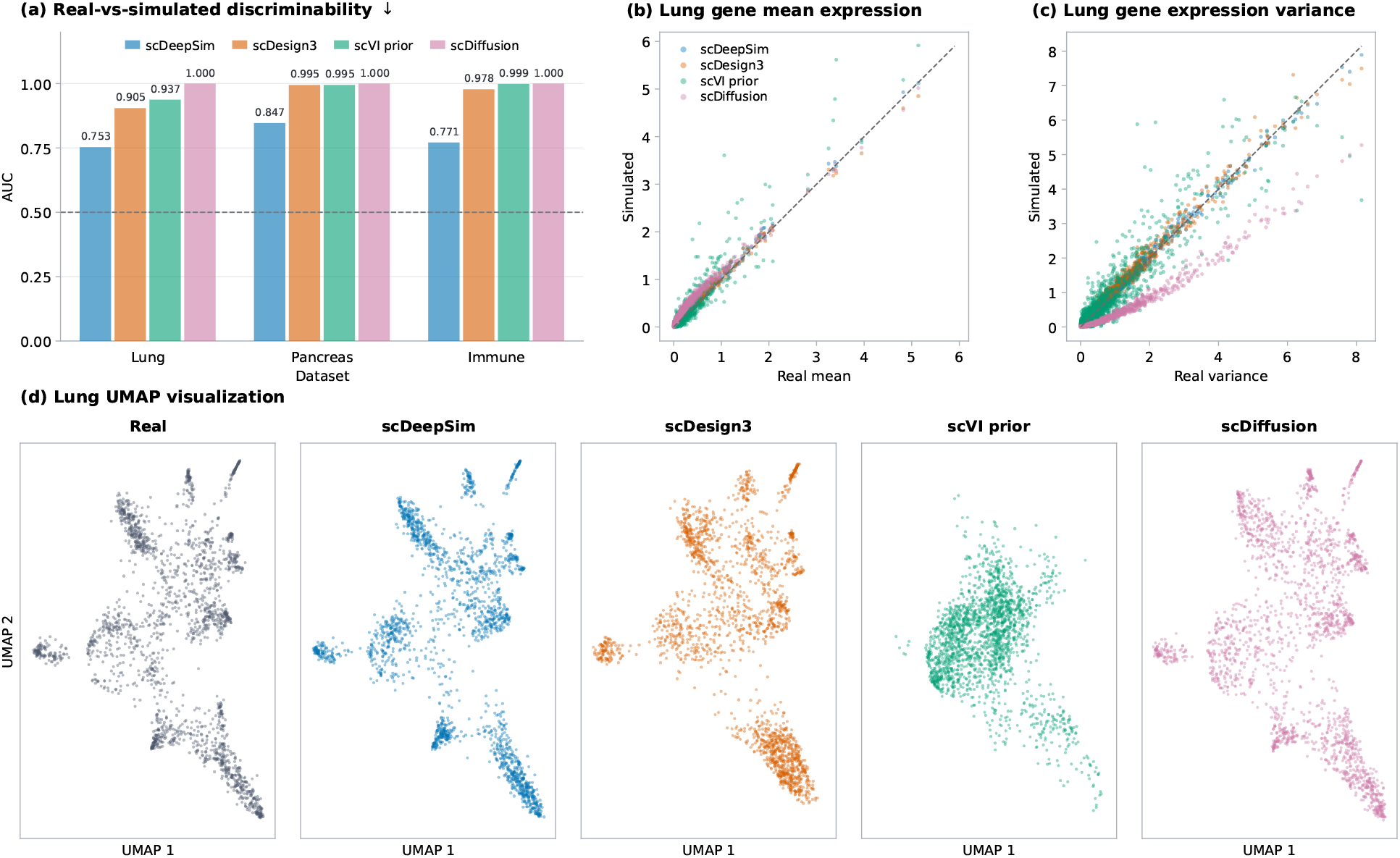
Simulation fidelity assessment across three scIB datasets, with gene-level and UMAP comparisons shown for lung. Within each dataset, all methods used the same training split and generated as many cells as the held-out reference. (a) Random forest ROC AUC for distinguishing real from simulated cells on lung, pancreas, and immune. The dashed line marks chance discrimination (AUC = 0.5); the downward arrow indicates that lower values are preferred. (b,c) Simulated versus real per-gene means and variances on lung, respectively; dashed lines indicate equality. (d) Joint UMAP projection of real and simulated lung cells. Colors identify methods throughout. Sampling dependencies and inclusion criteria for each dataset are described in Appendices B.3 and B.2.

**Fig. 3.**
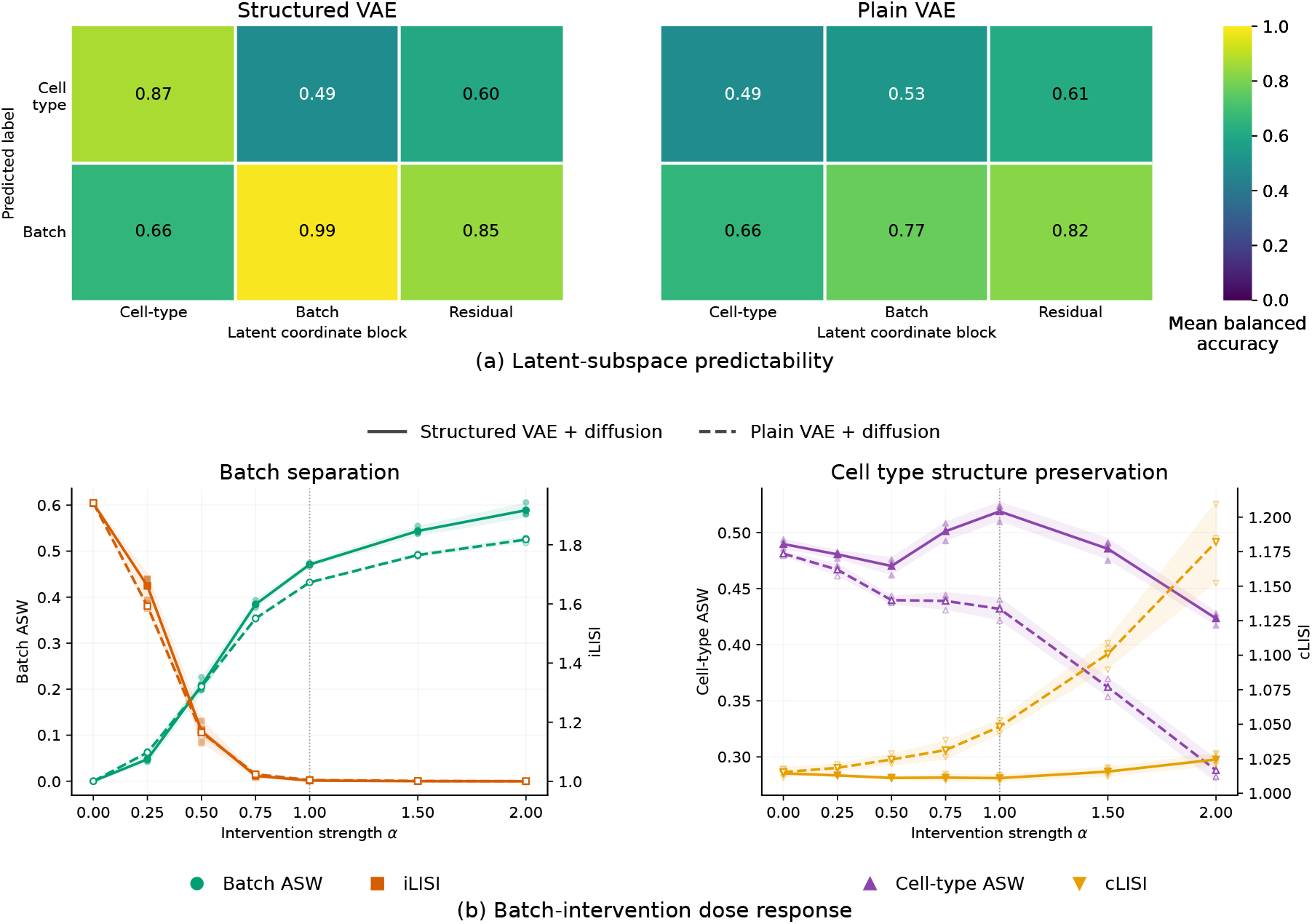
Latent organization and batch interventions on scIB human pancreas. (a) RF probe balanced accuracy for cell type and technology batch from the cell-type, batch, and residual subvectors. Separate heatmaps show structured and plain VAEs on the same color scale; colors and annotations show mean balanced accuracies across three training seeds. (b) Batch separation and cell-type preservation across intervention strength *α* for structured and plain VAE–diffusion models. The structured model changes its batch subspace; the plain model changes its full latent vector. The vertical dotted line marks *α* = 1; larger strengths extrapolate the fitted map. Lines and shading show means and *±*1 sample standard deviation across three training seeds; faint points show individual repetitions.

– We develop a generative model that learns structured single-cell representations and generates realistic samples from a learned population distribution.
– We introduce a post-hoc intervention strategy for varying selected technical and biological factors while largely preserving non-target structure, thereby providing tunable, intervention-defined ground truth.
– We demonstrate the utility of the simulator through applications to benchmarking batch integration and trajectory inference methods, showing how controlled simulations can reveal method-specific performance regimes as intervention strength and trajectory settings vary (Fig. 4 and 5).

**Fig. 4.**
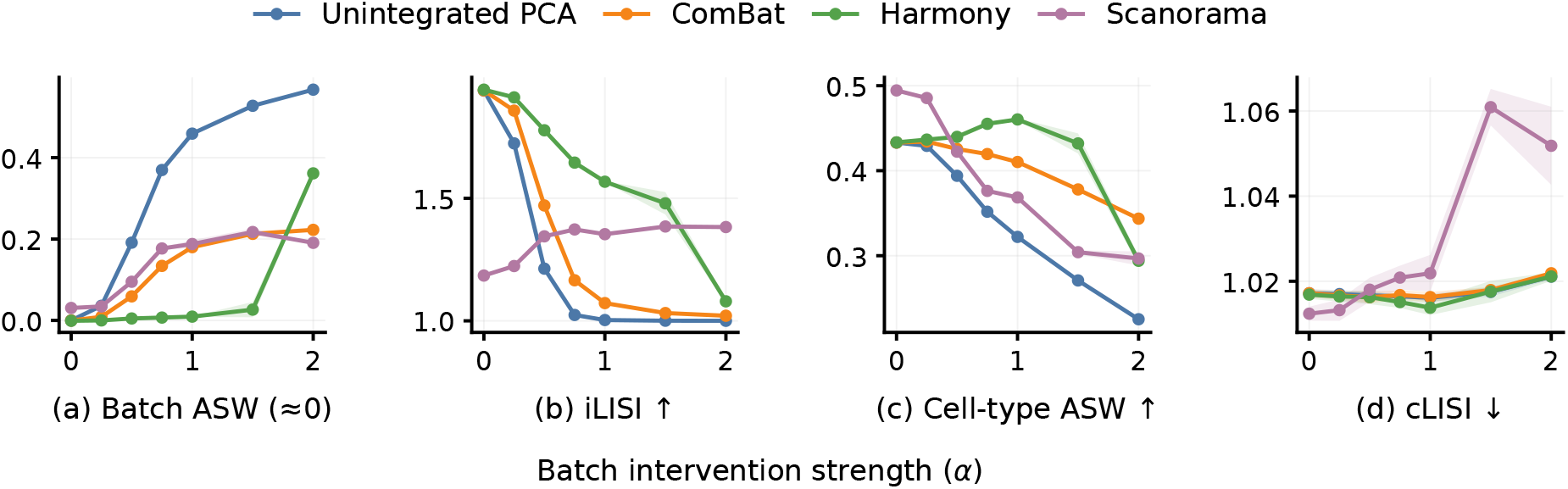
Batch integration across intervention strengths. *α* = 0 is the synthetic data without batch effect, *α* = 1 applies the fitted map, and *α >* 1 extrapolates it. Panels show (a) signed within-cell-type batch ASW, (b) iLISI, (c) cell-type ASW, and (d) cLISI in each method’s 30-dimensional representation of the combined cohorts. Lines and shading show means and *±*1 sample standard deviation across three generation seeds, conditional on the fixed simulator.

**Fig. 5.**
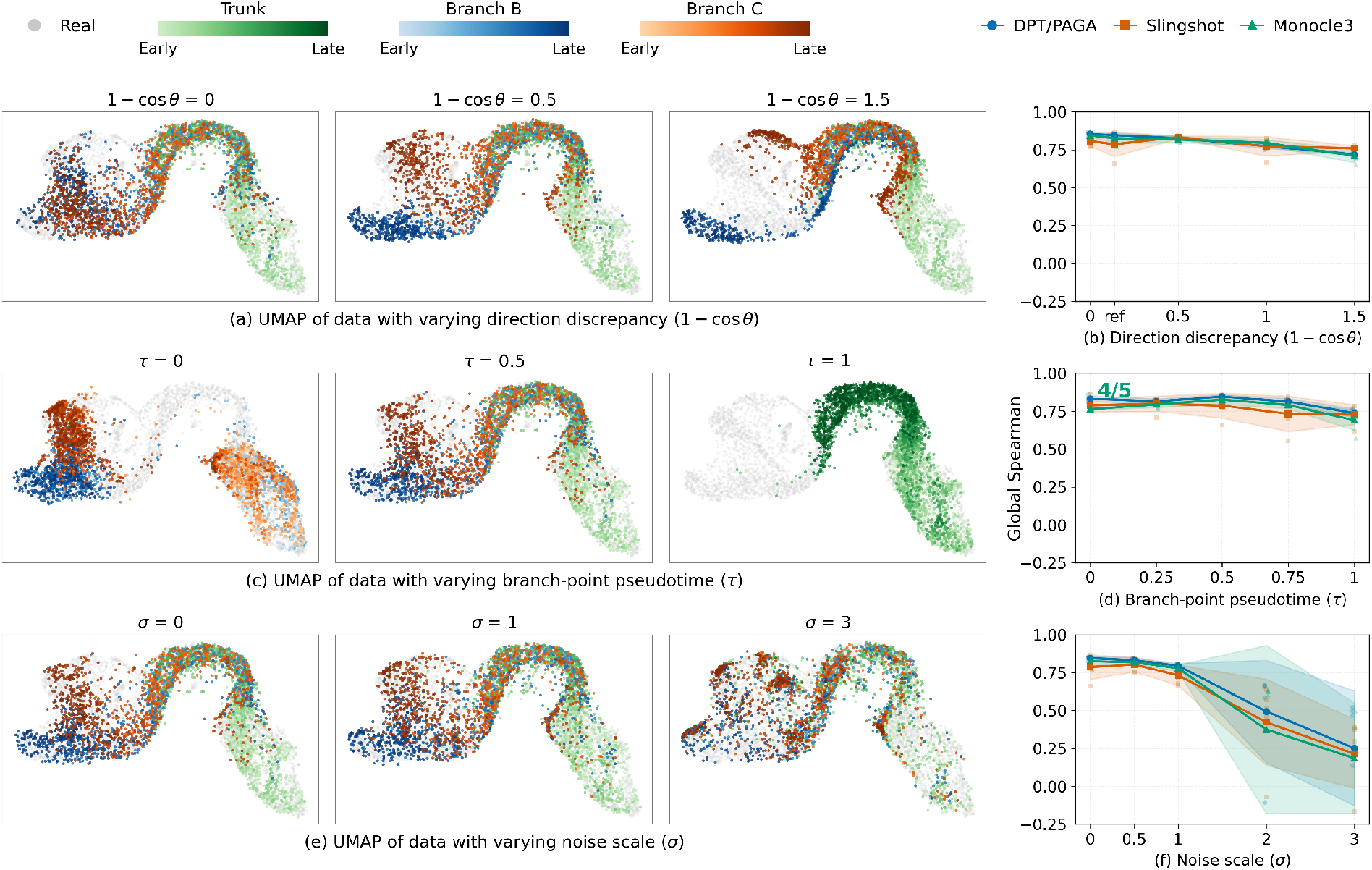
Controlled trajectory construction and TI benchmarking. (a,c,e) UMAP plots of data generated by varying direction discrepancy 1 −cos *θ*, branch-point pseudotime *τ*, and noise scale *σ*, respectively. Each group shows three representative settings: 1− cos *θ* ∈ {0, 0.5, 1.5}, *τ* ∈ {0, 0.5, 1}, and *σ* ∈ {0, 1, 3}. All datasets share the same projection learned from the real pancreas reference data (grey). Green, blue, and orange distinguish the trunk and two daughter branches; color intensity indicates ground-truth pseudotime within each lineage. (b,d,f) Corresponding global Spearman correlations for DPT/PAGA, Slingshot, and Monocle 3 across all five settings for each parameter. In (b), “ref” denotes the branch angle in the latent representation of the real dataset. Five replicate datasets were generated per setting using one trained model, and each dataset was evaluated by all three methods. Faint points show individual replicate correlations; lines and shaded regions show the mean and *±*1 sample standard deviation. Three replicate correlations below −0.25 fall outside the displayed range and remain included in the summaries. Monocle 3 returned non-finite pseudotime in one replicate at *τ* = 0, so the “4/5” annotation indicates that its summary uses four valid replicates out of five. All other summaries use five replicates.

## 2 Methods

We describe the structured representation and simulator (Section 2.1), the post-hoc interventions used to construct controlled datasets (Section 2.2), and the evaluation data, comparison methods, and metrics (Section 2.3).

### 2.1 Generative framework

Our goal is to generate new synthetic cells while allowing users to adjust selected sources of variation. For example, we retain batch variation in a designated part of the representation so that we can vary batch-effect strength while leaving the remaining latent representation unchanged.

To support this goal, scDeepSim uses two steps. First, it encodes factors of expression variation into separate latent subspaces on which we can then intervene. Second, it learns the joint distribution of these representations so that we can sample latent representations that do not correspond to any observed cell. We complete the first step with a structured zero-inflated truncated-normal VAE and the second with a conditional latent diffusion model.

#### Structured zero-inflated truncated-normal VAE

We use a structured VAE to encode cell expression profiles for three reasons: (i) to organize different sources of variation into distinct latent subspaces so that interventions can target individual factors without affecting the others; (ii) to allow manipulation of latent representations in Euclidean space while ensuring that decoded expression values remain realistic, whereas unrestricted interventions in raw expression space can produce negative values; and (iii) to support efficient diffusion training and sampling in a compact latent space [38, 37]. To achieve these goals, we combine a zero-inflated truncated-normal decoder with supervised and adversarial classifiers that guide the organization of the latent representation.

Let 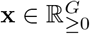 denote the log-normalized expression profile of a cell over *G* genes, obtained by normalizing counts in each cell to a total of 10^4^ and applying the natural-log transformation log(1 + *x*) element-wise. Let **z** ∈ ℝ^*d*^ denote its latent representation. Like scVI, we use a diagonal Gaussian encoder *q*_*ϕ*_(**z** |**x**) and standard Gaussian prior *p*(**z**) = *N* (**0**, *I*_*d*_) [22] for the representation [25].

The original scVI models raw counts with a zero-inflated negative binomial likelihood, conditioning its encoder and decoder on batch labels to account for batch variation [25]. However, its decoder also requires a cell-specific library-size factor or observed library size computed from the training data [42]. To avoid separately specifying or modeling library size during generation, we instead model log-normalized expression directly, producing profiles on the scale used for downstream benchmarking. The decoder outputs zero probabilities ***π***_*θ*_(**z**) ∈ (0, 1)^*G*^, locations **m**_*θ*_(**z**) ∈ ℝ^*G*^, and scales 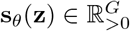. These define the zero-inflated truncated-normal distribution through the generative process:

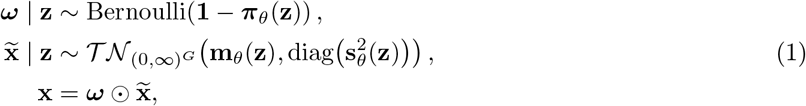

where ***ω*** and 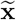 are conditionally independent, the Bernoulli coordinates are also conditionally independent, 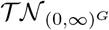 denotes a Gaussian distribution truncated to the positive orthant, and is element-wise multiplication. Thus, ***π***_*θ*_(**z**) controls the probability of zero expression, while the truncated-normal component models expression magnitude conditional on being positive. With this formulation, we can train the VAE by minimizing the standard negative evidence lower bound, ℒ_ELBO_ [22].

The ELBO alone does not assign factors to particular latent subvectors. One approach, used by scDisInFact, is to learn shared and condition-specific biological factors through separate encoders, while supplying batch labels directly to the decoder [57]. Condition classifiers and maximum mean discrepancy penalties guide the separation of these biological factors. In scDeepSim, we instead use a single encoder and explicitly tie biological and technical factors to a part of the representation on which we can intervene.

We partition the latent vector: 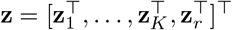, and train it so that **z**_*k*_ reflects an observed label *y*_*k*_ and **z**_*r*_ represents residual variation. In the experiments below, *K* = 2 and we write 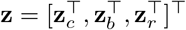, where **z**_*c*_, **z**_*b*_, and **z**_*r*_ correspond to cell type, batch, and residual variation, respectively, and *y*_*c*_ and *y*_*b*_ denote the observed cell-type and batch labels. To align the partition with the observed covariates, we train classifiers to predict each covariate from its own subvector. In our experiments, the supervised loss is

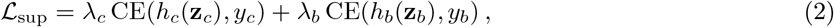

where *h*_*c*_ and *h*_*b*_ are neural network classifiers, CE is the cross-entropy loss, and *λ*_*c*_, *λ*_*b*_ control their contributions. These losses encourage cell type and batch to be predictable from their designated latent subvectors, but do not prevent their contributions to complementary coordinates. We therefore introduce conditional adversaries that attempt to predict each covariate from the complementary coordinates while conditioning on the other observed covariate:

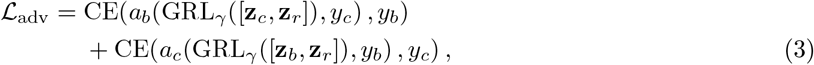

where *a*_*b*_ and *a*_*c*_ are conditional adversarial classifiers. Conditioning the batch adversary on cell type and the cell-type adversary on batch encourages *y*_*b*_ *⫫* (**z**_*c*_, | **z**_*r*_) *y*_*c*_ and *y*_*c*_ *⫫* (**z**_*b*_, | **z**_*r*_) *y*_*b*_, that is, it targets information about one factor that is not explained by the other. The gradient-reversal layer GRL_*γ*_ [13] leaves its input unchanged but multiplies encoder gradients by *−γ*. Hence the adversaries learn to predict the labels, while the encoder receives the reversed gradient and is trained to make *y*_*b*_ unpredictable from [**z**_*c*_, **z**_*r*_] conditional on *y*_*c*_, and *y*_*c*_ unpredictable from [**z**_*b*_, **z**_*r*_] conditional on *y*_*b*_.

The terms above address generation fidelity (*ℒ*_ELBO_), label prediction from assigned subvectors (*ℒ*_sup_), and reduced leakage (*ℒ*_adv_). Altogether, the VAE minimizes:

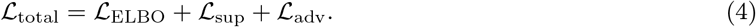

Section 3.2 compares how accurately each subvector predicts cell type and batch in structured and plain VAEs (Fig. 3a). Implementation details are provided in Appendix A.1.

#### Conditional latent diffusion

To improve sampling fidelity, we train a conditional diffusion model on the representations learned by the structured VAE, following latent diffusion approaches for image and single-cell generation [38, 32]. We use the DDPM framework [18] with classifier-free guidance [19]; categorical conditions select a learned subpopulation, such as a cell type or batch. Reverse diffusion generates *base latent vectors* that can be decoded directly or modified by the interventions described in Section 2.2. Implementation details are given in Appendix A.2, with numerical settings in Appendix B.4.

### 2.2 Post-hoc interventions

After sampling, we intervene on selected subvectors of the same base latent representations at different strengths. This produces matched synthetic datasets with known intervention parameters, which can be used to evaluate the performance of downstream computational methods under different known scenarios.

#### Factor-specific affine intervention

Consider an intervention on the subvector **z**_*k*_ assigned to a factor of interest. Let (***µ***_*k*,0_, ***Σ***_*k*,0_) and (***µ***_*k*,1_, ***Σ***_*k*,1_) denote the mean and covariance of the subvectors in source and target reference populations, respectively. For example, these populations can be cells from two different batches. Our goal is to transform the subvectors so that their mean and covariance match those in the target population. To do this, we define an affine map from the source to the target:

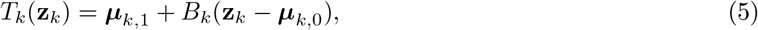

where *B*_*k*_ determines the variation of the transformed representations. We use 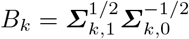 so that the transformed subvectors have a desired covariance. In practice, we use the sample means and covariances estimated from the source and target reference cells. More details are provided in Appendix A.3.

To control the strength of the intervention, we move each source subvector toward its mapped counterpart:

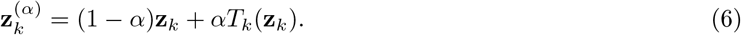

Here, 0 *≤ α ≤* 1 gives an interpolation, and *α >* 1 extrapolates beyond the target state. The intervention targets the selected factor by replacing only its designated subvector with 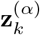, while leaving all other latent coordinates unchanged. The parameter *α* controls the strength of this targeted intervention.

#### Intervention-defined benchmark variables

We focus on batch effect and developmental trajectories as the primary factors to control. We use the affine intervention parameters to specify batch-effect strength and trajectory properties, providing known ground truth for benchmarking.

##### Batch-effect strength

For batch control, we estimate the moments of **z**_*b*_ from two reference batches, which serve as the source and target. Applying Equation 6 to **z**_*b*_ induces a batch effect while leaving the cell-type and residual subvectors unchanged.

##### Pseudotime

For a linear trajectory, we select labeled start and end states from the real data and estimate their moments using the non-batch subvectors 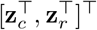. Applying the same affine intervention with *α* ∈ [0, 1] moves cells from the start to the end state while preserving **z**_*b*_. We assign *t* = *α* as the ground-truth pseudotime.

##### Branching trajectories

We select a start state, an intermediate waypoint, and two terminal states from the real data and use the same affine map (Equation 6) to construct a shared path from the start state to the waypoint and a branch from the waypoint to each terminal state. Joining the shared path to each branch gives two lineages. Each lineage has a shared trunk followed by one of the two branches, with pseudotime measuring progress along the joined path. The branching time *τ* ∈ [0, 1] is the pseudotime at which the paths split. Each branch has a mean direction from the waypoint to its terminal state. The angle *θ* between these directions controls how widely the branches diverge. Gaussian noise with scale *σ ≥* 0 controls the spread of cells around these paths in the non-batch latent subspace, and we keep the batch subvector fixed. Thus, pseudotime, lineage, branch angle, branching time, and noise scale are specified by construction. Details of path composition, pseudotime assignment, and noise injection are provided in Appendix A.3.

### 2.3 Evaluation methods

To assess whether scDeepSim generates realistic data with controllable variation for benchmarking, we evaluate it on human and mouse single-cell datasets. We assess fidelity using the human lung atlas of Vieira Braga et al. [49], with additional human pancreas [14, 34, 24, 43, 3, 54] and immune-cell datasets [36, 12, 47, 50, 1]. We use the processed releases assembled for the single-cell integration benchmarking (scIB) study [31]; these datasets also support evaluating latent organization and targeted interventions. The trajectory experiments use the mouse pancreatic endocrinogenesis data of Bastidas-Ponce et al. [4], obtained through scVelo [5]. Appendix B specifies dataset selection, numerical protocols, and evaluation conventions.

#### Simulation fidelity

We compare scDeepSim, scDiffusion [32], scVI [25], and scDesign3 [45] on the same cells, genes, and training/evaluation splits. Models are fitted to the training half, with internal validation confined to that half, and each model generates as many cells as the held-out reference.

Following classifier-based evaluations of single-cell simulation quality [33, 32], we assess fidelity by training a random forest to distinguish held-out real from simulated cells in log-normalized expression space. We report the area under the receiver operating characteristic curve (ROC AUC). An AUC near 0.5 indicates higher similarity between the real and simulated cells. We compare per-gene means and variances between real and simulated data using Pearson correlations. We also qualitatively compare population structure with joint UMAP visualizations.

Intervened datasets generally lack a real data counterpart against which fidelity can be assessed. As an indirect check, we follow the evaluation design of scGen [29]: we hold out one cell type from a target batch and train the VAE on the remaining cells. We estimate the batch effect intervention from these remaining cells and apply it to latent representations of the held-out cell type from the source batch. After decoding the transformed representations into synthetic expression profiles, we compare the gene-wise means and standard deviations of these profiles and the held-out target cells. Further details are provided in Appendix B.3.

#### Latent organization and targeted interventions

We compare structured and plain VAEs with matched architectures, training budgets, and cell splits. The plain VAE is trained without the supervised and adversarial objectives. We ask whether each label is easier to predict from its assigned subvector than from other subvectors. We train random-forest probes to predict cell type or batch from the latent subvector means E[**z**_*k*_| **x**] and report balanced accuracy on cells held out from probe training.

For each model, we independently generate two source cohorts. One cohort remains unchanged, and the other receives the batch effect intervention. We reuse the same base latent cells of the intervened cohort across a range of strengths *α*, applying a fixed batch-specific affine map (Section 2.2). The structured model changes only its batch subvector, and the plain model changes its full latent vector. Both models use diffusion-generated base cells. This comparison tests whether the structured VAE enables more targeted interventions when all other design choices are held fixed.

We measure separation between the two cohorts using within-cell-type batch average silhouette width (ASW) and the integration local inverse Simpson’s index (iLISI) [23]. Larger batch ASW and smaller iLISI indicate stronger cohort separation. Detailed definitions are provided in Appendix B.5. To assess biological separability within the intervened cohort, we measure cell-type ASW, the cell-type local inverse Simpson’s index (cLISI), and balanced accuracy for cell-type prediction with a random forest. If these separability measures remain stable as *α* increases, then the intervention preserves cell-type structure.

#### Downstream benchmark utility

We use scDeepSim to benchmark the performance of batch integration and trajectory inference methods under controlled simulation interventions.

For batch integration, we compare an unintegrated baseline, ComBat [21], Harmony [23], and Scanorama [17] on matched synthetic pancreas batches across intervention strengths *α*. We generate three replicate synthetic datasets at each intervention strength. All methods receive the same expression data and synthetic batch labels at each strength. We combine synthetic cells with no added batch effect and synthetic cells with an imposed batch effect, then apply each batch-integration method to this combined dataset. We evaluate whether the resulting representation mixes cells from the two batches while preserving cell-type distinctions.

Near zero within-cell-type batch ASW and higher iLISI indicate better batch mixing; higher cell-type ASW and lower cLISI indicate clearer cell-type structure.

For trajectory inference, we compare Monocle 3 [8], diffusion pseudotime (DPT) with partition-based graph abstraction (PAGA) in Scanpy [15, 52, 53], and Slingshot [46]. We train scDeepSim on the mouse pancreatic endocrinogenesis dataset provided by scVelo and use annotated cell states to construct bifurcating trajectories. We vary one trajectory parameter at a time: direction discrepancy 1 − cos *θ*, branching-point pseudotime *τ*, or noise scale *σ*, while holding the other two fixed. We generate five replicate datasets for each intervention setting and apply all three trajectory inference methods to the same datasets. We assess how well each method recovers the simulated ordering of cells by computing the global Spearman correlation between inferred pseudotime and the simulator’s ground-truth pseudotime across all cells. Values closer to one indicate more accurate recovery of this ordering. Detailed benchmark settings are provided in Appendix B.5.

## 3 Results

### 3.1 scDeepSim generates more realistic samples than established simulators

We first asked whether scDeepSim samples realistic cells from its learned distribution. We evaluated simulation fidelity on the human lung atlas of Vieira Braga et al. [49] and human pancreas and immune datasets, using the processed scIB releases [31]. Within each dataset, all methods used the same training split and generated as many cells as the held-out reference.

scDeepSim samples were harder to distinguish from held-out real cells than samples from the other methods across all three datasets (Fig. 2(a)). The random forest classifier’s ROC AUC was 0.753, 0.847, and 0.771 on lung, pancreas, and immune datasets, respectively, compared with 0.905, 0.995, and 0.978 for scDesign3 and 0.937, 0.995, and 0.999 for scVI prior sampling. scDiffusion had an AUC of 1.000 on all three datasets. Because values closer to 0.5 indicate that simulated cells are harder to distinguish from real cells, scDeepSim gave the closest match across all datasets. scDeepSim also closely reproduced the mean and variance of expression for individual genes. On the lung dataset, the simulated values lay near the identity line when plotted against those of held-out real cells (Fig. 2(b,c)). scDesign3 showed similarly close agreement, whereas scDiffusion systematically underestimated gene expression variance and scVI showed greater deviations in both means and variances. Similar results were observed in the pancreas and immune datasets, with the corresponding gene-level comparisons shown in Appendix C.1. In the UMAP embedding fitted jointly to real and simulated cells on the lung dataset, scDeepSim reproduced both the connected central structure and the smaller separated populations observed in the held-out real data (Fig. 2(d)).

Classifier choice also affects the fidelity assessment. In an auxiliary analysis of scDiffusion samples, KNN yielded a misleading near-chance AUC of 0.503 despite substantially reduced variability in the generated cells and perfect discrimination by a random forest on the same test cells (AUC 1.000, Appendix C.2). Assessing realism after intervention is challenging because real counterparts are generally unavailable for direct comparison. Nonetheless, validation against held-out real data provided indirect support for the realism of simulated cells after batch intervention, showing close agreement in gene-wise expression means and standard deviations (Appendix C.3).

### 3.2 Structured latent representations enable targeted batch interventions

We next asked whether the structured latent representations provided useful targets for intervention. We fitted scDeepSim on scIB human pancreas data. The cell-type and batch labels were most predictable from their designated cell-type and batch subvectors, respectively, and accuracy was higher than when using the corresponding subvectors from the plain VAE (Fig. 3(a)). The structured latent representations therefore provide useful intervention targets for controlling technical and biological variation. However, labels remained partly predictable from outside their designated subvectors. This leakage can result from the correlation between the observed batch and cell-type labels, for example when cell-type composition differs across batches.

We then applied the intervention defined by (6) to the batch subspace. As intervention strength *α* increased, within-cell-type batch ASW rose and iLISI fell, indicating that the intervention progressively increased batch separation as intended. In the structured model, cell-type ASW and cLISI remained comparatively stable over *α* ∈ [0, 1] (Fig. 3(b)). Thus, batch separation increased while cell types remained distinct and local neighborhoods retained their cell-type purity. The plain model also increased batch separation, but its cell-type ASW decreased and cLISI increased substantially. These changes indicate that its intervention also blurred cell-type distinctions and mixed different cell types within local neighborhoods. Under extrapolation (*α >* 1), cell-type ASW declined in both models, but the structured model better retained cell-type separation and neighborhood purity. This suggests that stronger interventions increasingly affect biological structure, although restricting the intervention to the batch subspace reduces this disruption. Together, these results show that scDeepSim provides more targeted control of batch variation while better preserving cell-type structure. Additional experiments on scIB immune and lung showed similar patterns (Appendix C.4).

### 3.3 Controlled simulations reveal how data conditions shape method performance

#### Batch integration across intervention strengths

We next asked whether controlling batch intervention strength could affect batch-integration method rankings. Using the same model trained on the scIB human pancreas dataset, we generated synthetic batches for each *α*. We compared an unintegrated PCA baseline, ComBat, Harmony, and Scanorama on how well they integrate the synthetic batches, whose batch effects have varying strengths, and the unintervened synthetic batch across three random seeds (Fig. 4).

As intervention strength increased, unintegrated cells became more separated by batch. At the weakest nonzero intervention, the methods performed comparably. Harmony’s advantage in batch mixing and cell-type separation became more pronounced as *α* increased toward 1. It retained the highest mean iLISI through *α* = 1.5, but its mixing deteriorated at *α* = 2. At that strongest intervention, Scanorama had the highest mean iLISI, but also had higher cLISI, indicating a trade-off between batch mixing and cell-type structure preservation. This is consistent with the trade-off between batch removal and biological conservation reported by Luecken et al. [31]. ComBat had the highest cell-type ASW at that strength, but weaker batch mixing, as measured by iLISI. Thus, varying intervention strength revealed different responses in batch mixing and cell-type structure. In the unintervened *α* = 0 data, Scanorama reduced iLISI relative to unintegrated PCA, indicating poorer local batch mixing after correction. This is consistent with previous reports that batch correction can reduce batch mixing even when no batch effect is present [30, 2]. Appendix C.5 reports means and sample standard deviations across three replicates for batch ASW, iLISI, cell-type ASW, and cLISI for each method at *α* ∈ {0, 1, 2}.

#### Controlled trajectory interventions reveal sensitivity to data conditions

Finally, we used controlled trajectory interventions to compare inference methods and examine how data conditions affect pseudotime recovery. We varied branch angle *θ*, branch-point pseudotime *τ*, and noise scale *σ* separately, evaluating three trajectory-inference (TI) methods against the simulator’s ground-truth pseudotime using global Spearman correlation. Increasing noise led to the largest decrease in pseudotime correlations across the tested trajectory settings (Fig. 5(f)). With branch geometry fixed, mean correlations across the three TI methods fell from 0.786–0.846 at *σ* = 0 to 0.185–0.251 at *σ* = 3. Increasing noise also produced greater variation among generation replicates, and recovery became less accurate as lineage-specific structure mixed in the fixed UMAP projection (Fig. 5(e)). Increasing branch angle made the daughter branches appear more separated in the UMAP (Fig. 5(a)).

However, the three methods showed similar mean global Spearman correlations across the tested branch angles and branch-point pseudotimes (Fig. 5(b,d)). Large variation across replicates in the high noise settings also made differences between method means less clear. These experiments also show how controlled simulations can reveal which data conditions make pseudotime harder to recover. Appendix C.5 reports means and sample standard deviations of global Spearman correlation for each method at selected branch angles, branch-point pseudotimes, and noise levels.

## 4 Discussion

Even when ground truth is available, benchmarking on a real dataset establishes performance under the observed conditions, leaving open how that performance would change if the data were different. Our results show that deep generative simulation can extend such benchmarks by combining realistic sampling with continuous control of selected factors. In the batch-integration study, varying intervention strength around the fitted batch transformation (*α* = 1) revealed how method performance and the trade-off between batch mixing and cell-type preservation changed under weaker or stronger batch effects. Controlled trajectory simulations likewise showed that greater separation between branches did not necessarily improve recovery of temporal ordering. These findings reinforce the importance of assessing method robustness across data conditions, as emphasized in recent work on single-cell benchmarking [10]. By varying individual properties, scDeepSim allows users and method developers to examine when performance deteriorates and which trade-offs emerge, providing a practical framework for controlled stress testing.

Several limitations remain. Batch and cell-type information is still partly encoded outside their assigned latent subspaces, and stronger interventions can weaken cell-type structure (Section 3.2). Thus, changing one subspace does not guarantee that other properties remain unchanged. Current controls also do not support gene-specific interventions. Our validation against held-out data (Appendix C.3) provides evidence that manipulated samples can reproduce the overall structure and selected expression statistics of held-out data. However, interventions beyond the observed range may move latent representations into regions poorly covered by the training data, where decoding may produce less realistic expression profiles. The lack of direct experimental counterparts for these samples also makes their realism difficult to assess.

To improve the realism of manipulated samples, future work could pretrain scDeepSim on larger, more diverse cell atlases or adapt existing foundation models, such as scGPT and scFoundation [11, 16]. Exposure to a broader range of cellular states could help the model decode a broader region of latent space reliably, providing better support for interventions beyond the range observed in an individual dataset. Incorporating gene-program information, as in VEGA and expiMap [44, 28], may enable controls over program-associated expression. Our auxiliary KNN experiment (Appendix C.2) also showed that nearly constant classifier predictions can yield near-chance AUC despite detectable distributional differences, motivating further examination of classifier-based protocols for evaluating simulation quality. Increasing noise may obscure the ordering information in the data, limiting how accurately any method can recover pseudotime. Future work could also investigate whether this imposes a theoretical upper bound on recovery accuracy and how that bound changes with noise level.

## A Method details

### A.1 Structured VAE details

The VAE uses fully connected encoder and decoder networks without residual connections. Decoder heads output the location, a softplus-transformed positive scale, and a logit for the zero probability in Equation (1). Reconstruction loss sums the gene-wise negative log-likelihoods and averages over cells. Supervised classifiers use their assigned contiguous latent subvectors; each conditional adversary receives the complementary coordinates together with an embedding of the other covariate’s label. Supervised cross-entropy weights (Equation (2)) and gradient-reversal strength (Equation (3)) apply separately: the adversarial classifiers minimize unweighted cross-entropy, while gradient reversal acts on their latent inputs.

Posterior means are used for the predictability probes, whereas posterior samples supply the diffusion training data. The plain VAE ablation retains the backbone with supervised and adversarial heads removed. Network dimensions, loss weights, and dataset-specific partitions are listed in Appendix B.4.

### A.2 Latent diffusion details

After VAE training, we encode each training cell once using a reparameterized posterior sample **z**_0_ *∼ q*_*ϕ*_(**z** |**x**). A residual MLP U-Net learns the latent distribution using the DDPM formulation [18] and *v*-prediction [41]. Writing *ā*_*t*_ for the cumulative signal-retention factor of the cosine noise schedule [35], the training input and target are

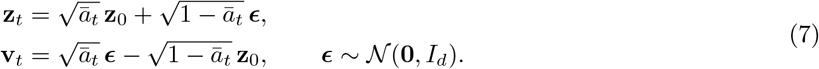

The denoiser **v**_*ψ*_ is trained to predict this target by minimizing

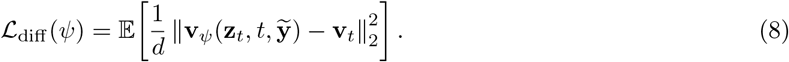

The expectation is over training latent vectors and their labels, uniformly sampled timesteps, Gaussian noise, and label dropout. Here, 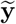 denotes the categorical condition after dropout: its embedding is randomly replaced by a learned null embedding so that the same network learns conditional and unconditional predictions. The objective is mean squared error averaged over latent coordinates and cells, with equal weight across timesteps.

At generation time, classifier-free guidance combines conditional and unconditional predictions to control the influence of the requested labels [19]. We use an exponential moving average (EMA) of the denoiser weights for sampling, as in DDPM [18]. Starting from standard Gaussian noise, full-step DDPM sampling produces new latent vectors, which are passed to the ZITN decoder, optionally after intervention. Our diffusion and guidance implementation is adapted from `denoising-diffusion-pytorch` [51]. Numerical training, guidance, and sampling hyperparameters are given in Table 3.

### A.3 Post-hoc intervention details

#### Whitening–recoloring

For the endpoint map in Equation (5), we use

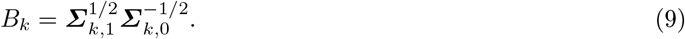

The square roots are computed from the sample covariance matrices estimated from the reference data, and the source square root is inverted. For nonsingular source covariance, this map whitens source variation and recolors it with the target covariance, giving *T*_*k*_(**z**_*k*_) target mean ***µ***_*k*,1_ and covariance ***Σ***_*k*,1_ when the input has the specified source moments. Batch reference populations and moment estimation are specified in Appendix B.

#### Trajectory composition and pseudotime

Let *A* denote the initial state, *W* a shared waypoint, and *B, C* the terminal states. All trajectory maps act on 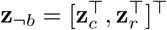 and preserve **z**_*b*_. For 0 *< τ <* 1, affine interpolation forms a trunk from *A* to *W* and two daughter branches from *W* to *B* and *C*. We divide the cells at the trunk endpoint between the two daughter branches. These cells initialize the branch paths at the shared branch point. The angle *θ* between the two waypoint-to-terminal mean vectors defines branch geometry.

The local interpolation coordinate *α* is mapped to common pseudotime by

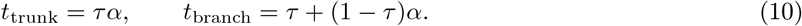

For interior values of *τ*, this changes branch timing without changing segment paths or endpoints. At each sampled common time, the trunk receives twice as many cells as each daughter branch, keeping the total cell count per time point constant. With a uniform time grid, trunk occupancy is approximately proportional to *τ*, and each daughter’s occupancy to (1 *− τ*)/2. At *τ* = 0, the construction uses direct *A → B* and *A → C* maps with no trunk; at *τ* = 1, it contains only the *A → W* trunk with no daughter branches.

#### Trajectory noise

After interpolation, we add noise only in the non-batch subspace:

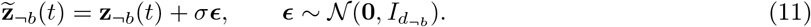

The noise scale controls dispersion around the latent paths without changing the assigned pseudotime, lineage labels, or reference endpoint definitions.

## B Experimental details

### B.1 Data and code availability

Code implementing the scDeepSim method and manuscript experiments is available at https://github.com/LangtianM/scDeepSim. All datasets are publicly available. The human lung, pancreas, and immune datasets are obtained from the processed scIB releases [31]; the mouse pancreatic endocrinogenesis dataset is provided through scVelo [4, 5]. The auxiliary KNN diagnostic uses Tabula Muris data [48].

### B.2 Datasets and preprocessing

Table 1 lists the selected populations. For human lung, pancreas, and immune cells, we use the processed scIB releases [31]. The original lung data were generated by Vieira Braga et al. [49]; the release combines 10x Genomics and Drop-seq profiles from that study (Drop-seq accession GSE130148).

**Table 1.**
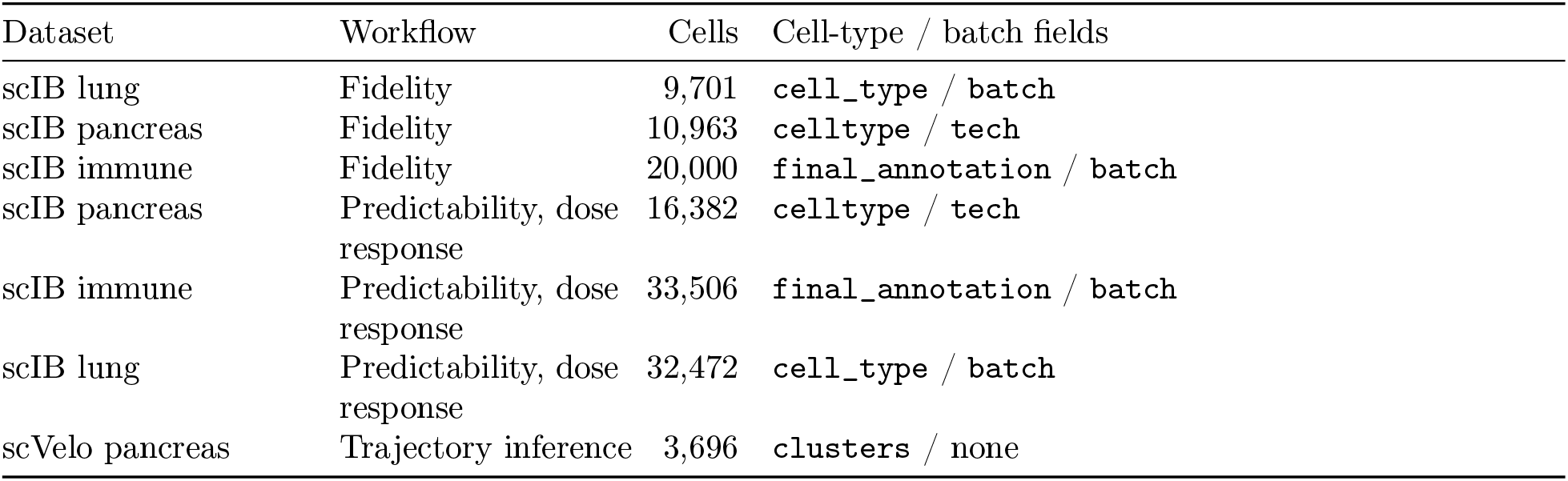
Dataset selection. Counts are cells retained for each workflow; fidelity counts precede the outer training/evaluation split.

The human pancreas release combines six studies: Grün et al. [14] (GSE81076, CEL-seq), Muraro et al. [34] (GSE85241, CEL-seq2), Lawlor et al. [24] (GSE86469, Fluidigm C1), Segerstolpe et al. [43] (E-MTAB-5061, SMART-seq2), Baron et al. [3] (GSE84133, inDrop), and Xin et al. [54] (GSE81608, SMARTer).

The human immune release combines bone marrow profiles from Oetjen et al. [36] (GSE120221) with peripheral blood profiles from Freytag et al. [12] (GSE115189), Sun et al. [47] (GSE128066), Villani et al. [50] (GSE94820), and the 10x Genomics 10k PBMC healthy-donor release with v3 chemistry [1].

The trajectory data originate from Bastidas-Ponce et al. [4]; we use the processed pancreatic endocrino-genesis dataset supplied by scVelo 0.3.4 with its `clusters` labels.

All workflows retain cells with at least 10 detected genes and genes detected in at least two cells, select 2,500 highly variable genes (HVGs) using Seurat-v3 selection, normalize each cell’s selected-gene total to 10,000, and apply log(1 + *x*). Fidelity additionally requires finite, nonnegative, integer-like counts, cell-type labels, and at least six cells per label.

### B.3 Experimental protocols

#### Simulation fidelity

Single-cell simulators use different ways to estimate parameters and assume different levels of conditioning context, so their fidelity is not always directly comparable [9]. We group sampling procedures by the information used to generate each synthetic cell:

– *De novo sampling* draws *x∼ p*_*ψ*_(*x*| *c*), where the generative mechanism and its key parameters *ψ* are predefined or specified by the user. Here, *c* denotes conditions or design variables, such as cell type, batch, trajectory structure, or perturbation. For example, dyngen generates cells from a predefined or user-designed regulatory backbone and kinetic model [7].
– *Learned-distribution sampling* draws *x ∼p*_*θ*_(*x*| *c*), where *θ* is fitted to real training data and a new *x* is sampled from the fitted distribution. For example, scDesign3 fits marginal and copula models to real data and then samples from the fitted distribution [45].
– *Observation-conditioned sampling* draws *x∼ p*_*θ*_(*x* |*x*_obs_, *c*), where a particular observed profile *x*_obs_ determines the cell-specific representation used at sampling time and *θ* is fitted from training data. For example, scVI posterior sampling takes real cells as input and generates a new expression profile conditioned on each input cell [25].

Observation-conditioned procedures have access to cell-specific information that learned-distribution procedures do not, so fidelity values should be compared within sampling categories. Our main fidelity experiment therefore compares learned-distribution methods fitted to the same training split and producing the same number of samples. We apply the same evaluation separately to observation-conditioned procedures to assess how well they reproduce real-data distributions when conditioned on observed cells (Figs. 8–10).

We now describe the learned-distribution sampling methods used in our experiments:

– *scDeepSim*. scDeepSim fits a ZITN-VAE to the training profiles and a diffusion model to the resulting latent states. At sampling time, reverse diffusion generates new latent states under the requested conditions, and the VAE decoder samples the corresponding expression profiles with the latent states.
– *scDiffusion*. scDiffusion trains an autoencoder followed by a latent diffusion model. New latent states are drawn through reverse diffusion and decoded into expression profiles, placing this procedure in the learned-distribution category.
– *scDesign3*. scDesign3 fits negative-binomial marginal models and a Gaussian copula to the training counts, conditional on cell type, and samples new counts from the fitted distribution.
– *scVI prior sampling*. scVI does not provide a direct prior-sampling interface for this use. We therefore draw latent vectors manually from the standard Gaussian prior and pass them to the fitted scVI generative module. We also sample library-size and covariate inputs from training cells as sampling inputs.

The following are the observation-conditioned sampling methods we evaluated for diagnostic purposes:

– *ZINB-WaVE*. ZINB-WaVE fits a zero-inflated negative-binomial model with observed covariates and learns sample-level latent factors. Its simulation procedure uses the same latent factors of the training data, so every output remains conditioned on a training cell.
– *scVI posterior sampling*. Training cells are sampled with replacement to match the evaluation sample size. For each sampled cell, scVI uses the observed counts to infer a cell-specific latent representation, and generates one new count profile while retaining the input cell’s total-count scale.
– *scDeepSim VAE posterior sampling*. To evaluate the ZITN-VAE separately from the diffusion model, we also use its posterior sampling procedure. For each evaluation profile, the VAE infers a cell-specific latent representation, samples from that representation, and decodes one reconstructed expression profile.

#### Experimental configuration

We first remove cells whose supplied `counts` layer contains non-finite or negative values, or values farther than 10^−6^ from the nearest integer. We then apply the cell, gene, and label filters described above. We split cells equally into outer training and evaluation sets, stratified by cell type, using seed 42. This gives 4,850/4,851 training/evaluation cells for lung, 5,481/5,482 for pancreas, and 10,000/10,000 for immune. Each dataset has one model fit per method; scDeepSim’s internal 80/20 validation split is confined to the outer training half. All methods use the same selected genes and outer split, receiving counts or log-normalized expression as required. Generated sample sizes equal the evaluation counts. The original scDesign3 study selected 1,000 highly variable genes to reduce computational cost [45]. Here, we fit marginals for all 2,500 genes but restrict the Gaussian copula to the 1,000 genes with the highest variance in raw counts to keep computation tractable. Observation-conditioned diagnostics use the inputs for each method described above.

##### Validation against held-out data

To indirectly assess whether the data remain realistic after intervention, we followed the evaluation design in scGen [29]. We removed one cell type from the target batch during training and learned how the batches differed using the other cell types. We then transformed cells of the removed type from the source batch and compared their predicted gene expression with the real cells held out from the target batch. We used scIB pancreas, with the `inDrop3` batch as the source and the `smartseq2` batch as the target. In six separate experiments, we withheld all cells of one type from the `smartseq2` batch. Each target group was excluded before gene selection and VAE fitting. We selected 2,500 highly variable genes from the remaining cells, normalized selected-gene totals to 10,000, and applied log1p to both the remaining and withheld populations. After excluding the held-out target cells, we used 80% of the remaining cells to fit the VAE and reserved 20% for validation during training. Each VAE used the shared settings in Appendix B.4. Seeds 42–44 gave three independent fits per holdout.

For each holdout, we fitted the map using cells from the other five selected cell types. Within each type, we sampled equal numbers of source and target cells, taking the smaller available count; seed 42 fixed these memberships across training repetitions. We pooled the matched populations and estimated the means and covariances of their batch subvectors from posterior samples. We then applied the affine map in Equations (5) and (9) at *α* = 1 to encoded real source cells of the withheld type. Only the batch subvector changed; cell-type and residual coordinates remained fixed. Predicted target expression was sampled from the VAE decoder.

For each fit, we calculated *R*^2^ separately for gene-wise means and standard deviations of log-expression across all 2,500 selected genes, using the held-out real target summaries as observed values and the corresponding simulated summaries as predictions.

##### Structured latent representations and targeted batch interventions

Three training seeds (42–44) define structured/plain VAE pairs on the same pancreas cells, with one diffusion fit for each of the six VAEs. Within each pair, the backbone initialization and unstratified 80/20 model split are shared; diffusion reuses that split. Random-forest probes use separate 80/20 splits stratified by the target label, shared across models and latent subspaces within each seed. These are within-dataset representation probes; their test cells need not have been excluded from VAE training.

Targeted interventions use the `inDrop3` batch as the source and the `smartseq2` batch as the target. Each independently generated source cohort contains 500 cells per type (3,000 cells per cohort). For each fitted model, source/target reference moments and the batch-specific affine map are estimated from real-cell latent representations and frozen. The unchanged cohort and the base latents of the intervened cohort are reused for *α* ∈ {0, 0.25, 0.5, 0.75, 1, 1.5, 2}. Section 2.2 defines the transformation.

The additional immune and lung experiments use the same paired training, probe, and intervention design, with three training seeds per dataset. They use the supplied counts layer without the fidelity workflow’s integer-count filter or immune subsampling, yielding the populations in Table 1. Each generated cohort contains 500 cells per included cell type. Source and target reference populations are matched by cell type to fit the whitening-recoloring map; included types have at least two cells in each reference batch. Table 2 lists the directions and cohort sizes. The immune cohorts contain CD14+ monocytes, CD16+ monocytes, CD20+ B cells, CD4+ T cells, CD8+ T cells, megakaryocyte progenitors, NK cells, NKT cells, and plasmacytoid dendritic cells. The lung cohorts contain B cell, Ciliated, Dendritic cell, Endothelium, Fibroblast, Lymphatic, Macrophage, Neutrophil_CD14_high, Secretory, T/NK cell, and Type 2 populations.

**Table 2.** Batch-intervention directions and generated composition. Each cohort has 500 cells per included cell type; each dataset uses three structured/plain training pairs.

| Dataset | Source batch | Target batch | Cell types | Cells per cohort |
|---|---|---|---|---|
| Pancreas | inDrop3 | smartseq2 | 6 | 3,000 |
| Immune | 10X | Freytag | 9 | 4,500 |
| Lung | 4 | 5 | 11 | 5,500 |

##### Downstream benchmarking

###### Batch integration

One structured VAE–diffusion pair trained with seed 42 and its saved whitening-recoloring map from the `inDrop3` batch to the `smartseq2` batch are fixed throughout the benchmark. Sampling seeds 42–44 define three independent pairs of source-conditioned cohorts, each with 500 cells per type for acinar, activated stellate, alpha, beta, delta, and ductal cells. Each combined dataset has 6,000 cells and the same 2,500 genes used by the source model. Within each repetition, the unchanged cohort and the base latents of the intervened cohort are reused across the seven strengths above. Only the intervened cohort’s 32-dimensional batch subspace changes; the decoder seed is reset before each strength. The model and map are reused without retraining or selection by integration performance.

All methods receive identical decoded expression and synthetic batch labels for each repetition and strength, without further normalization, scaling, or gene selection. Unintegrated PCA, Harmony, and Scanorama share the initial 30-dimensional PCA representation; ComBat corrects expression before PCA. ComBat uses no additional covariates, and cell-type labels are used only for evaluation. Scanorama uses exact matching with 20 neighbors, smoothing parameter 15, and alignment cutoff 0.1. PCA and Harmony random states are fixed at 42. The implementations use Scanpy 1.11.5, harmonypy 0.0.10, and Scanorama 1.7.4. All 84 method–strength–repetition evaluations completed with finite metrics.

###### Trajectory inference

One VAE–diffusion pair is trained at seed 42; generation seeds 42–46 define five repetitions. Ductal, Ngn3 high EP, Beta, and Alpha serve as the start, waypoint, and two terminal anchors. Each generation seed supplies a 916-cell Ductal latent pool. Sampling 100 cells at each of 21 time values for each of two terminal trajectories gives 4,200 cells per setting and repetition. The direction grid is 1 − cos *θ* ∈ {0, ref, 0.5, 1, 1.5} at *τ* = 0.5 and *σ* = 0; the branch-time grid is *τ* ∈ {0, 0.25, 0.5, 0.75}, 1 at the reference direction and *σ* = 0; the noise grid is *σ* ∈ {0, 0.5, 1, 2, 3} at the reference direction and *τ* = 0.5. The reference direction is determined by the anchor means in the fitted model’s latent space. Generation seeds are paired across settings and methods. The cell with minimum simulated pseudotime supplies the common root.

### B.4 Model and training settings

Table 3 specifies the scDeepSim hyperparameters used in the experiments. Algorithmic details are provided in Appendix A.

**Table 3.** Selected scDeepSim model, training, and sampling settings. For datasets without batch labels, the latent partition is 32 cell-type and 96 residual coordinates, with no batch head or conditional adversaries. The plain ablation retains the same backbone and training budget. Complete settings and implementation details are available in the scDeepSim repository (https://github.com/LangtianM/scDeepSim), including the experiment configurations in `experiments/configs/`.

| Setting | Value |
|---|---|
| *VAE* |  |
| Encoder / decoder widths | (512, 256) / (256, 512) |
| Latent dimension | 128: cell type / batch / residual = 32 / 32 / 64 |
| KL weight | 1 |
| Each supervised CE weight | 5 |
| Gradient-reversal strength | 5 |
| *Latent diffusion* |  |
| Denoiser | Residual MLP U-Net, widths (512, 256, 256, 128) |
| Objective / noise schedule | *v*-prediction / cosine [35] |
| Categorical conditions | Cell type, jointly with batch when available |
| Sampler / steps | Full-step DDPM; 1,000 sampling steps |
| Label dropout probability | 0.1 |
| Guidance scale / rescaling blend | 1.5 / 0.7 |
| *Training* |  |
| Optimizer | AdamW [26] |
| VAE / diffusion learning rate | 10<sup>−3</sup> / 10<sup>−4</sup> |
| Epochs / batch size | 200 per stage / 256 |
| Model validation | Seeded, 80/20 cell split, reused across stages |
| Latent representations | Posterior means for random forest probes; one posterior sample per cell for diffusion training |

### B.5 Evaluation conventions

#### Simulation fidelity and latent probes

Real-versus-simulated classification uses full log-normalized expression, 100 random-forest trees with unrestricted depth, and an 80/20 classifier split using seed 42. ROC AUC is evaluated on the classifier holdout. Gene-moment agreement uses Pearson correlations across selected genes. Latent probes use 100 trees with maximum depth 10 and the splits stratified by target label described above. Balanced accuracy is mean recall across target classes [6].

#### Batch and cell-type structure

All dose-response ASW and local-diversity calculations use a fixed 30-dimensional PCA representation fitted to each dataset’s real expression matrix. Batch metrics use the combination of unintervened and intervened data; cell-type metrics use intervened data only. For batch ASW, Euclidean silhouettes [39] use the intervention-defined batch labels separately within each cell type. For cell *i* of type *t*, let *a*_*it*_ be its mean distance to other cells of the same batch and type, and let *b*_*it*_ be the smallest mean distance to cells of another batch within that type. With *n*_*t*_ cells in type *t* and *T* cell types, we average silhouettes equally across types:

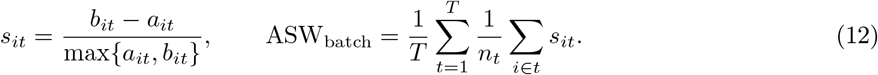

Cell-type ASW uses cell-type labels instead of batch labels and averages silhouettes across all cells without stratification.

We use local inverse-Simpson diversity, following the label-diversity interpretation of LISI [23], with 30 equally weighted nearest neighbors excluding self. For the fraction *p*_*ic*_ of label *c* in cell *i*’s neighborhood,

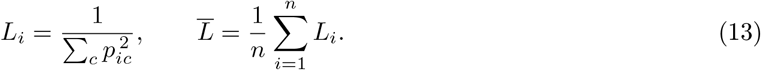

Here iLISI uses intervention-defined batch labels, with larger values indicating more mixing; cLISI uses cell-type labels, with smaller values indicating purer neighborhoods. These raw scores use neither distance/perplexity weighting nor unit-interval rescaling.

Dose-response balanced accuracy uses full decoded expression in the intervened synthetic data. At each strength, a random forest with 100 trees and maximum depth 10 is refitted on an 80/20 split at seed 42. This measures cell-type separability within each intervened population.

#### Batch correction benchmark

For batch integration, all four ASW and LISI metrics instead use each method’s 30-dimensional representation of the original and intervened cohorts. Batch ASW still weights cell types equally, and LISI still uses raw scores calculated from 30 nearest neighbors. Predictability and dose-response summaries report means and sample standard deviations across three training seeds; batch-integration summaries use one trained simulator with three different random seeds for generation. Sample standard deviations use denominator *n −* 1.

#### Trajectory inference benchmark

Global Spearman correlation compares the complete simulated time axis with native Slingshot average pseudotime (`slingAvgPseudotime`), native Monocle 3 pseudotime, or Scanpy DPT pseudotime in the DPT/PAGA workflow. Predictions are aligned by cell identifier. A run is invalid if pseudotime is incomplete, non-finite, or otherwise cannot yield a finite all-cell correlation. Summaries report means and standard deviations across valid generation repetitions and their valid counts out of five.

## C Supplementary experimental results

### C.1 Simulation fidelity

In each figure in this subsection, the top row shows a joint UMAP projection of real and simulated cells. The bottom row shows random forest ROC AUC and simulated versus real per-gene means and variances, from left to right. The dashed line in the AUC panel marks chance discrimination (AUC = 0.5); the diagonal dashed lines in the gene-level panels indicate equality.

#### Learned-distribution sampling

On scIB pancreas and immune data, scDeepSim produced samples that were harder to distinguish from held-out real cells than the other learned-distribution methods, while closely matching gene-wise means and variances (Figs. 6 and 7).

**Fig. 6.**
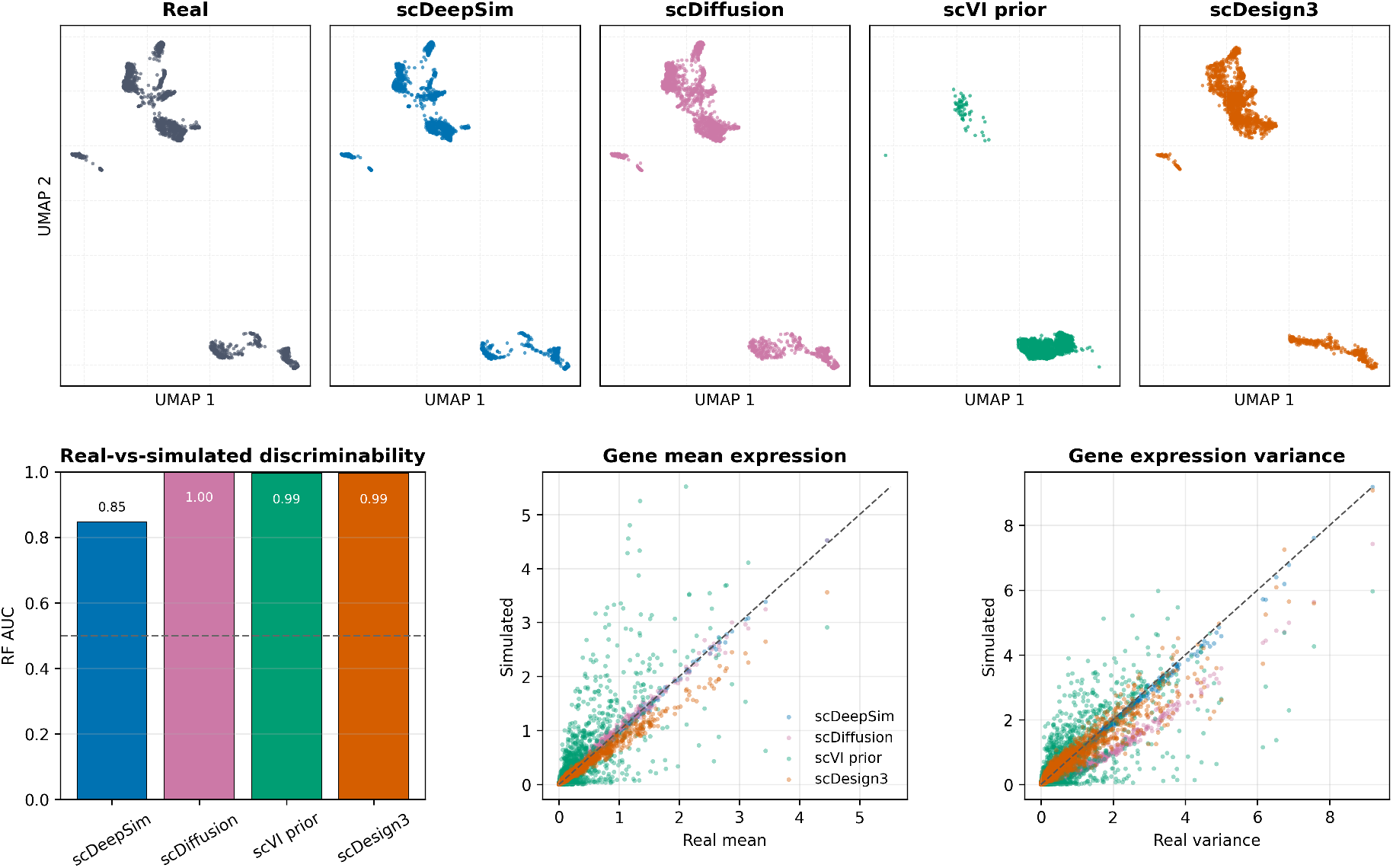
Learned-distribution simulation fidelity on scIB pancreas.

**Fig. 7.**
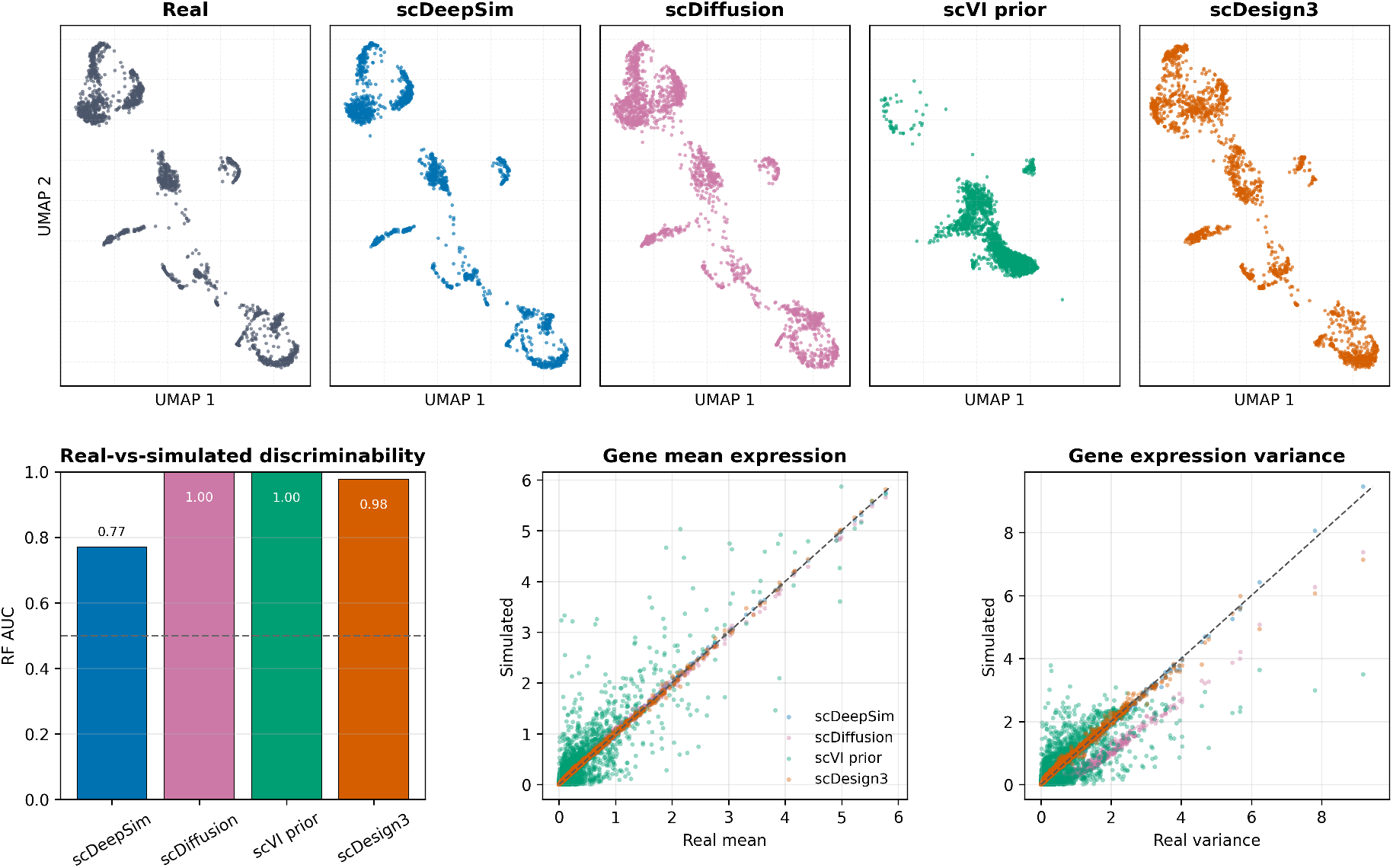
Learned-distribution simulation fidelity on scIB immune.

#### Observation-conditioned sampling

Gene-wise means and variances were closely reproduced across all three datasets, while discriminability differed by method and dataset (Figs. 8–10). scVI posterior samples had the lowest real-versus-simulated AUC on lung and immune; scDeepSim VAE reconstruction and scVI posterior sampling had similar AUC on pancreas, with a slightly lower value for the VAE.

**Fig. 8.**
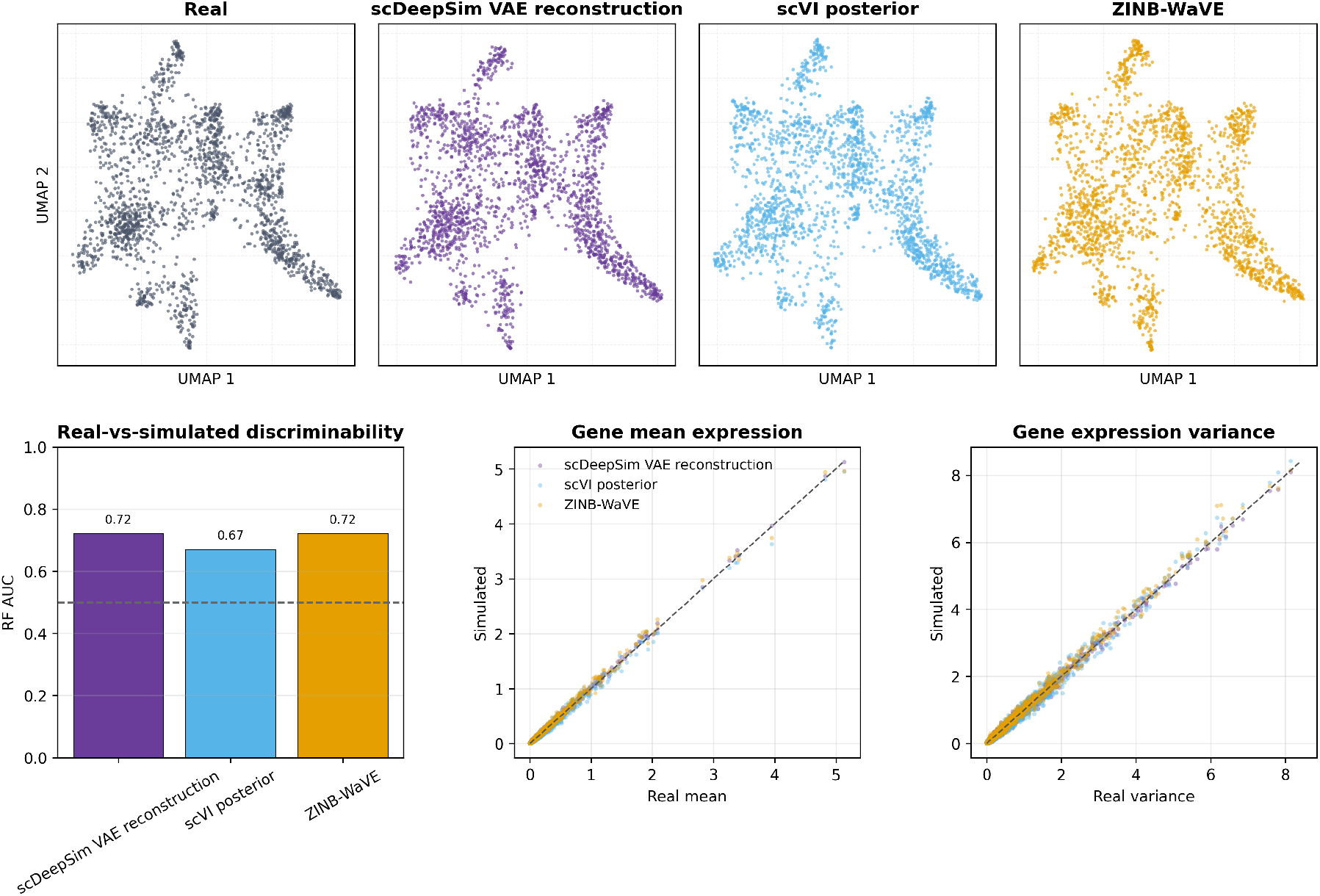
Observation-conditioned simulation fidelity on scIB lung.

**Fig. 9.**
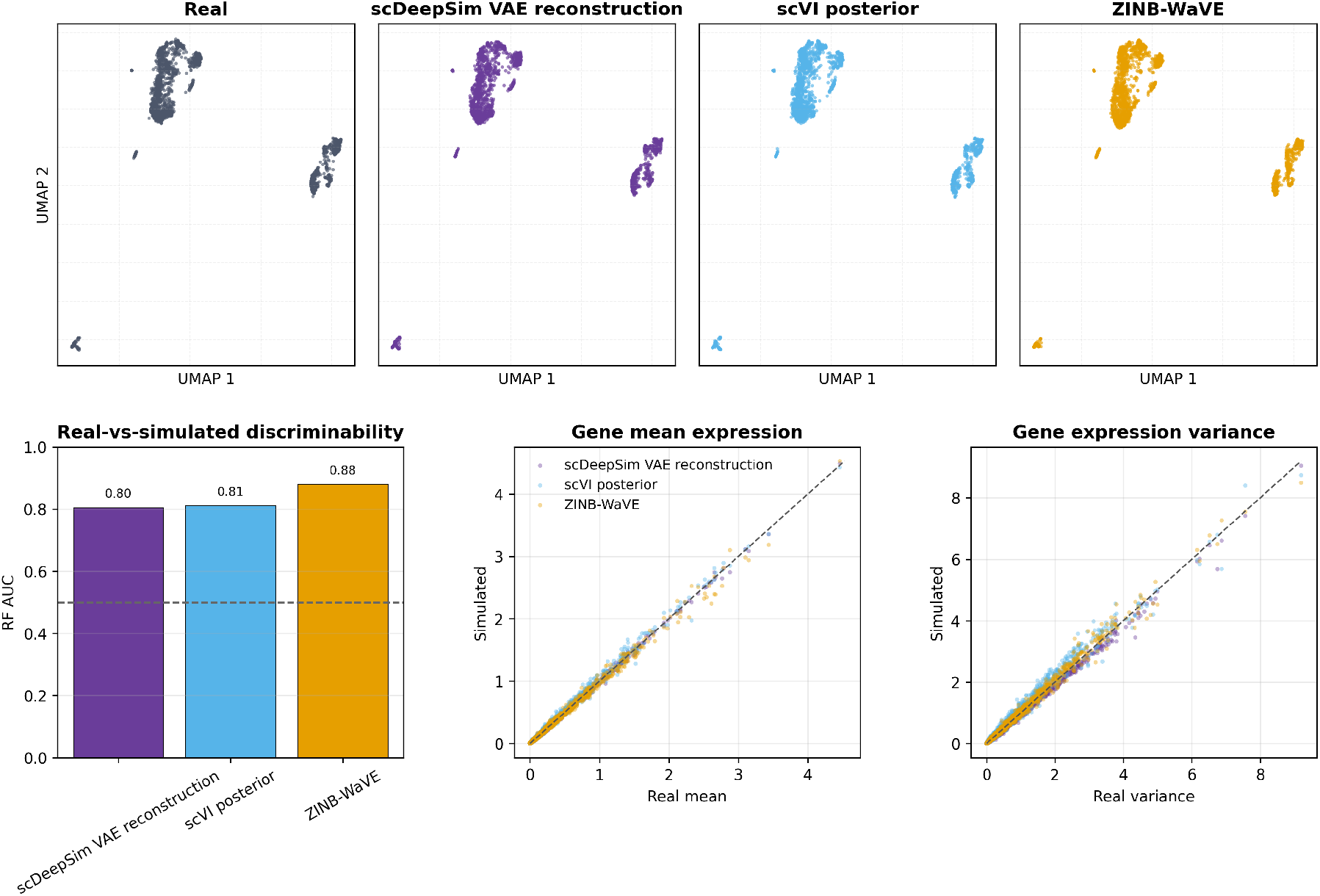
Observation-conditioned simulation fidelity on scIB pancreas.

**Fig. 10.**
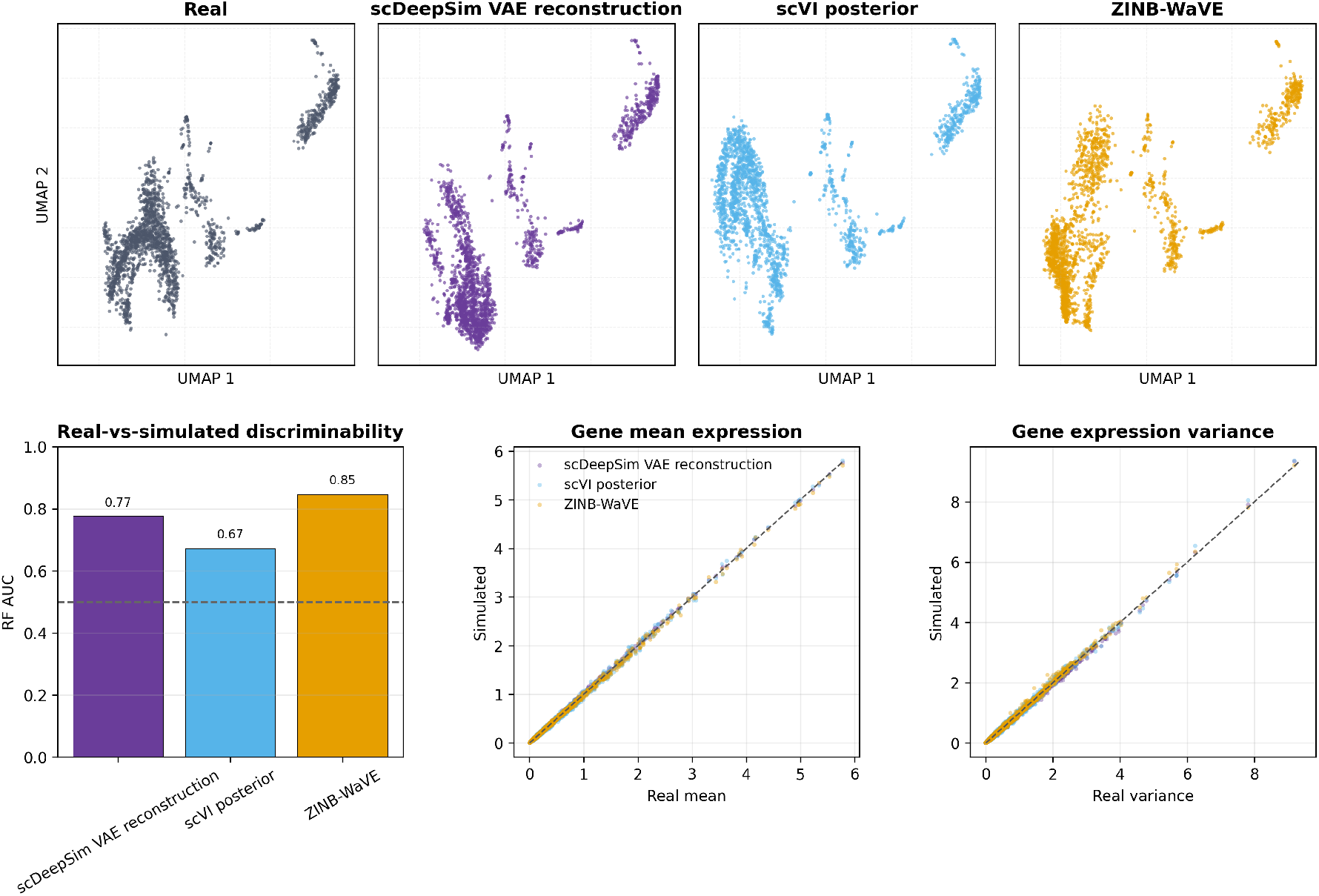
Observation-conditioned simulation fidelity on scIB immune.

### C.2 Classifier sensitivity in real-versus-simulated evaluation

The scDiffusion study reports near-chance *k*-nearest-neighbor (KNN) AUC for conditionally generated cells evaluated within each cell type [32]. To examine the interpretation of such scores, we used the Tabula Muris dataset [48], following the dataset choice in the scDiffusion evaluation. We fitted scDiffusion on the dataset and generated simulated cells with unconditional sampling following their documented procedure. We performed a separate diagnostic using 1,000 real cells and 1,000 simulated cells. Following the KNN evaluation settings in scDiffusion’s released evaluation notebook, ^4^ we used all 18,996 genes in normalized log1p expression space, five Euclidean neighbors, uniform weights, and a 70/30 train–test split with random seed 1. We also trained a random forest using 100 trees with unrestricted depth and seed 42 to evaluate the same cells.

KNN achieved an AUC of 0.503 but classified all 600 test cells as generated. The predicted probability of being real was zero for 598 cells and 0.2 for the other two, both of which were real. The near-chance AUC therefore arose from almost entirely tied scores. In contrast, the random forest achieved an AUC of 1.000 on the same test cells. Therefore, the near-chance KNN AUC alone does not necessarily indicate indistinguishable data distributions.

The lower variance of the generated cells helps explain why the KNN predictions collapsed. The sum of per-gene variances was 724 for generated cells, compared with 2,269 for real cells, and their mean profiles were close: the squared distance between them was 5.51. The generated cells were therefore more tightly concentrated around a similar mean. In this setting, even real test cells tended to be closer to more generated training cells than to real ones. As a result, 598 of the 600 test cells had only generated cells among their five nearest neighbors, giving the zero predicted probability for being real.

### C.3 Validation against held-out data

The validation against held-out data described in Appendix B.3 produced gene-expression summaries across six withheld cell-type–batch populations. Figure 11 illustrates the result of transferring alpha cells from the `inDrop3` batch to the `smartseq2` batch for seed 42. Across cell types, prediction *R*^2^ values are shown in Table 4.

**Table 4.** Agreement in gene-wise means and standard deviations between simulated and held-out real data across all 2,500 selected genes. Entries are mean*±* sample SD of *R*^2^ across three training seeds (42–44). Gene-SD denotes the gene-wise expression standard deviation being predicted; the uncertainty reports variation in *R*^2^ across seeds.

| Held-out cell type | Gene-mean prediction *R*<sup>2</sup> | Gene-SD prediction *R*<sup>2</sup> |
|---|---|---|
| Acinar | 0.863 ± 0.021 | 0.839 ± 0.008 |
| Activated stellate | 0.788 ± 0.013 | 0.682 ± 0.030 |
| Alpha | 0.949 ± 0.002 | 0.919 ± 0.006 |
| Beta | 0.933 ± 0.003 | 0.914 ± 0.003 |
| Delta | 0.743 ± 0.014 | 0.701 ± 0.014 |
| Ductal | 0.894 ± 0.023 | 0.856 ± 0.015 |

**Fig. 11.**
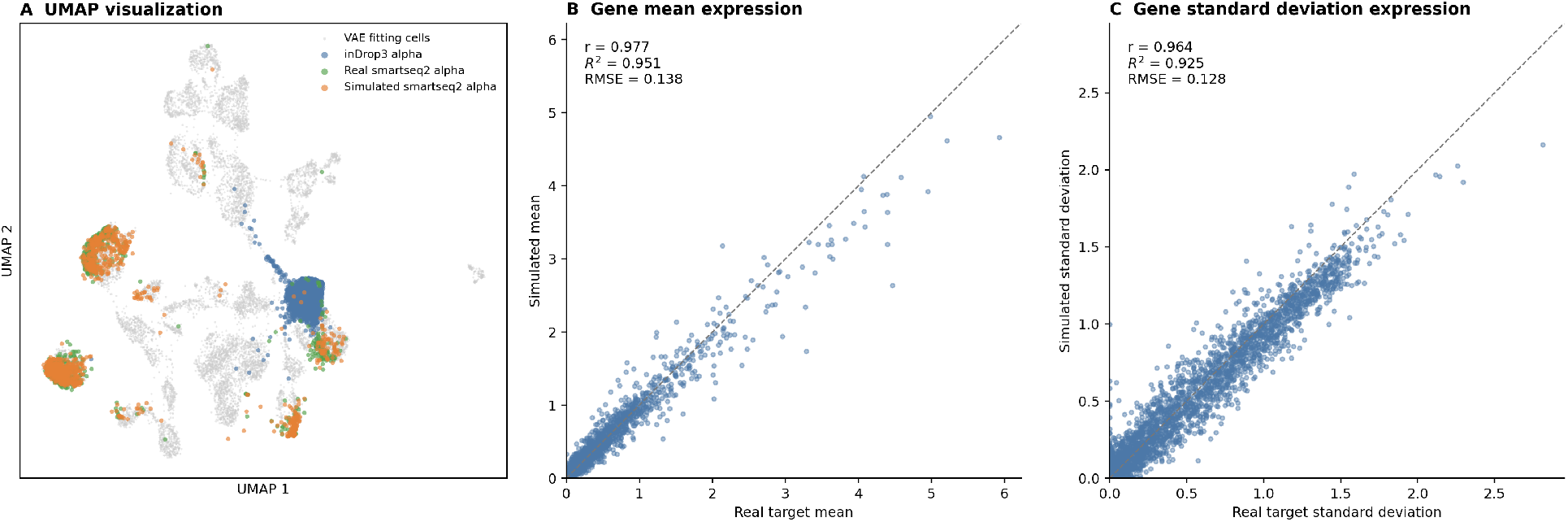
Illustrative held-out alpha-cell transfer from the `inDrop3` batch to the `smartseq2` batch, using seed 42 and whitening–recoloring at *α* = 1. Predictions condition on encoded real source cells, followed by batch-subvector intervention and VAE decoding. (A) PCA/UMAP fitted on VAE training cells (gray), with real source (blue), held-out real target (green), and predicted target (orange) alpha cells projected into the same embedding. (B–C) Predicted versus real target gene means and standard deviations across 2,500 selected genes in log-normalized expression space. Dashed lines indicate identity.

Agreement in these expression summaries supports the ability of scDeepSim to synthesize realistic intervened expression patterns.

### C.4 Structured latent representations and targeted batch interventions

On scIB immune and lung datasets, the designated subspaces were the most predictive of their respective labels, and increasing intervention strength produced stronger batch separation (Figs. 12 and 13). The structured model better retained cell-type structure than the plain model. Preservation weakened with increasing strength on immune. On lung, the structured model’s cell-type ASW and cLISI remained comparatively stable through *α* = 1, with preservation weakening under extrapolation.

**Fig. 12.**
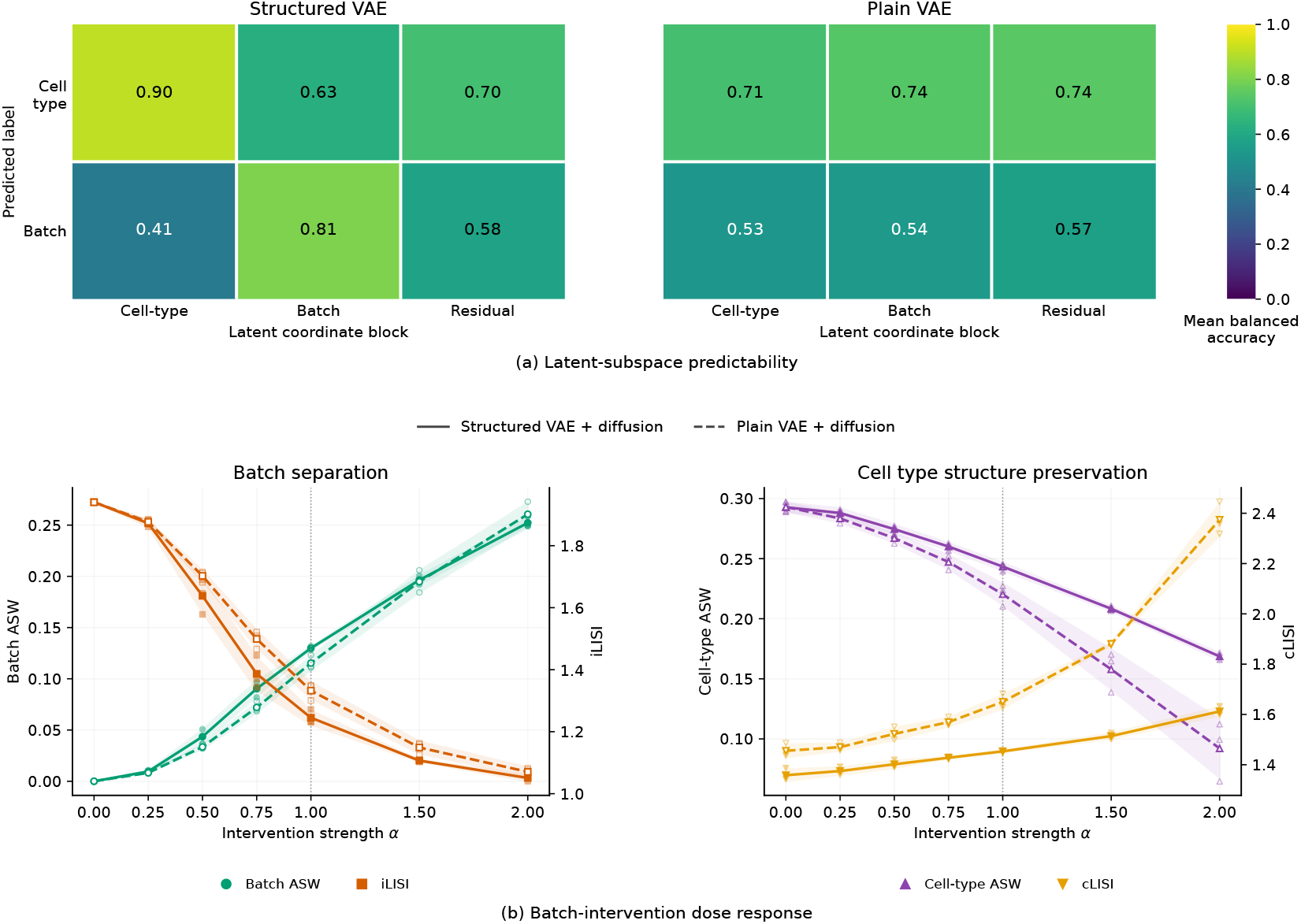
Latent organization and batch interventions on scIB immune, shown as in Fig. 3.

**Fig. 13.**
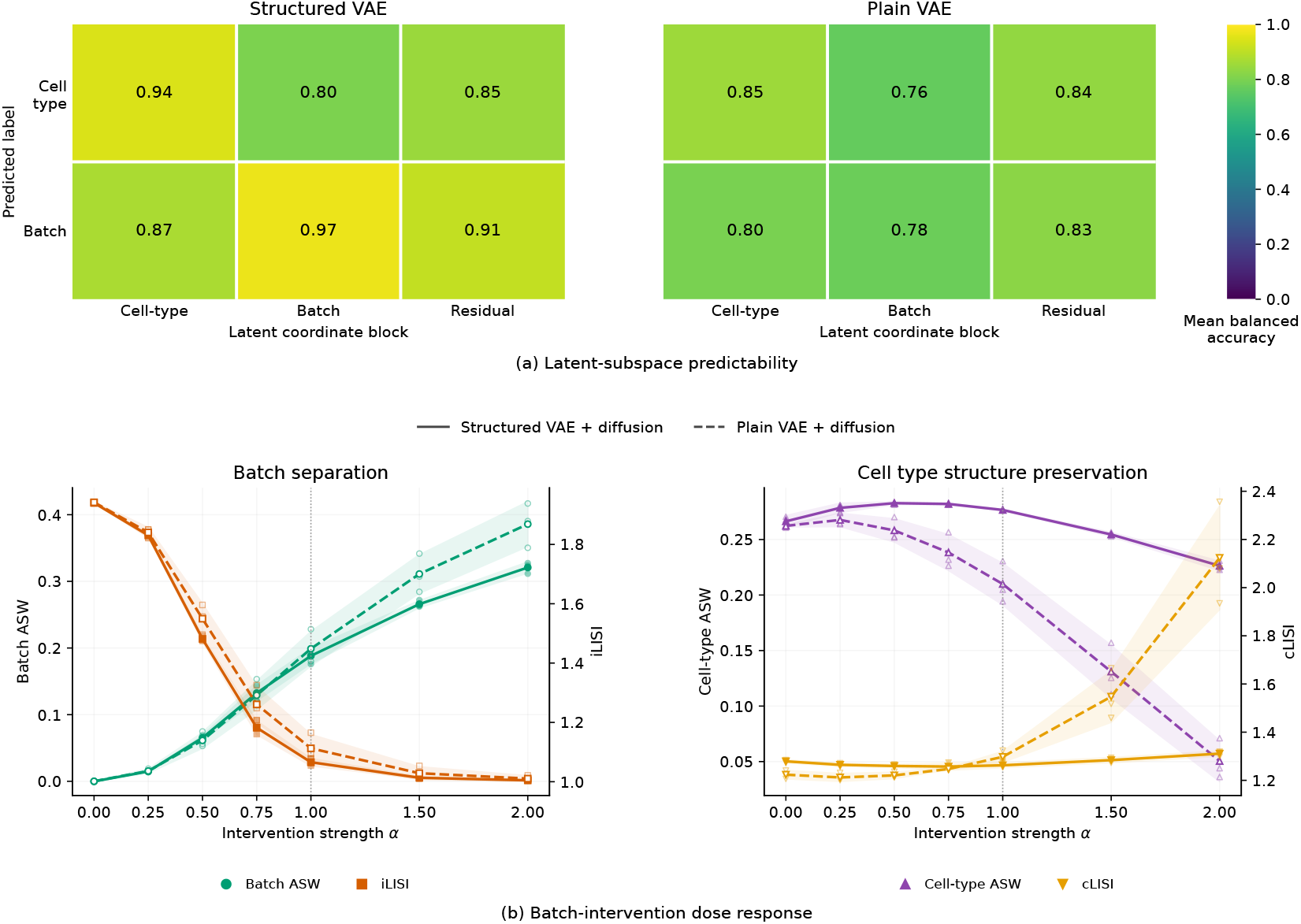
Latent organization and batch interventions on scIB lung, shown as in Fig. 3.

### C.5 Downstream benchmarking

#### Batch integration

Table 5 reports the synthetic null, fitted-map, and strongest extrapolation settings. Figure 4 shows the response curves across all seven intervention strengths.

**Table 5.** Batch-integration metrics at selected intervention strengths. Values are mean *±* sample standard deviation across three generated datasets, conditional on the fixed seed-42 simulator and map. Batch ASW near zero and higher iLISI indicate successful batch effect correction; higher cell-type ASW and lower cLISI indicate stronger cell-type signal preservation.

| α | Method | Batch ASW | iLISI | Cell-type ASW | cLISI |
|---|---|---|---|---|---|
| 0 | Unintegrated PCA | 0.0000 ± 0.0001 | 1.9394 ± 0.0028 | 0.4331 ± 0.0013 | 1.0172 ± 0.0009 |
|  | ComBat | −0.0002 ± 0.0001 | 1.9401 ± 0.0027 | 0.4331 ± 0.0013 | 1.0172 ± 0.0010 |
|  | Harmony | −0.0009 ± 0.0001 | 1.9431 ± 0.0028 | 0.4333 ± 0.0013 | 1.0169 ± 0.0010 |
|  | Scanorama | 0.0306 ± 0.0004 | 1.1847 ± 0.0014 | 0.4945 ± 0.0017 | 1.0124 ± 0.0016 |
| 1 | Unintegrated PCA | 0.4591 ± 0.0004 | 1.0032 ± 0.0003 | 0.3222 ± 0.0020 | 1.0161 ± 0.0009 |
|  | ComBat | 0.1801 ± 0.0008 | 1.0724 ± 0.0017 | 0.4103 ± 0.0018 | 1.0163 ± 0.0012 |
|  | Harmony | 0.0093 ± 0.0005 | 1.5675 ± 0.0025 | 0.4603 ± 0.0013 | 1.0138 ± 0.0017 |
|  | Scanorama | 0.1878 ± 0.0095 | 1.3536 ± 0.0025 | 0.3686 ± 0.0010 | 1.0219 ± 0.0043 |
| 2 | Unintegrated PCA | 0.5672 ± 0.0017 | 1.0002 ± 0.0001 | 0.2253 ± 0.0019 | 1.0211 ± 0.0006 |
|  | ComBat | 0.2226 ± 0.0011 | 1.0208 ± 0.0004 | 0.3436 ± 0.0025 | 1.0219 ± 0.0003 |
|  | Harmony | 0.3614 ± 0.0086 | 1.0796 ± 0.0082 | 0.2940 ± 0.0053 | 1.0211 ± 0.0010 |
|  | Scanorama | 0.1907 ± 0.0016 | 1.3825 ± 0.0029 | 0.2967 ± 0.0082 | 1.0518 ± 0.0092 |

#### Trajectory inference

Table 6 reports selected settings from Figure 5. Of the 225 method runs, 224 returned finite pseudotime for every cell and yielded a finite global Spearman correlation. Monocle 3 returned non-finite pseudotime in one replicate at *τ* = 0, leaving four valid replicates for that summary. All other method–setting combinations had five valid replicates.

**Table 6.** Global Spearman correlation at selected trajectory settings. Five replicate datasets were generated per setting using one trained model, and each dataset was evaluated by all three methods. Values show the mean *±* sample standard deviation, with the number of valid replicates included in parentheses. We varied one setting at a time, keeping the others at the reference branch angle, branch-point pseudotime *τ* = 0.5, or noise scale *σ* = 0, as applicable.

| Axis | Setting | DPT/PAGA | Slingshot | Monocle 3 |
|---|---|---|---|---|
| Branch angle | 60° | 0.826 ± 0.013 (5) | 0.833 ± 0.010 (5) | 0.817 ± 0.020 (5) |
|  | 90° | 0.774 ± 0.014 (5) | 0.772 ± 0.064 (5) | 0.796 ± 0.020 (5) |
|  | 120° | 0.719 ± 0.008 (5) | 0.760 ± 0.017 (5) | 0.710 ± 0.044 (5) |
| Branch time | 0 | 0.832 ± 0.020 (5) | 0.788 ± 0.044 (5) | 0.762 ± 0.004 (4) |
|  | 0.5 | 0.846 ± 0.013 (5) | 0.786 ± 0.081 (5) | 0.826 ± 0.031 (5) |
|  | 1 | 0.737 ± 0.028 (5) | 0.727 ± 0.065 (5) | 0.691 ± 0.067 (5) |
| Noise | 0 | 0.846 ± 0.013 (5) | 0.786 ± 0.081 (5) | 0.826 ± 0.031 (5) |
|  | 2 | 0.494 ± 0.338 (5) | 0.423 ± 0.284 (5) | 0.375 ± 0.555 (5) |
|  | 3 | 0.251 ± 0.378 (5) | 0.215 ± 0.228 (5) | 0.185 ± 0.365 (5) |

## Footnotes

4 https://github.com/EperLuo/scDiffusion/blob/d04430ed27f76e0e94b2399767868971e15b6020/exp_script/script_static_eval.ipynb

